# A coronavirus protease activates sensory neurons via TRPA1 to drive sneeze, cough, and pain

**DOI:** 10.1101/2024.01.10.575114

**Authors:** Sonali S. Mali, Zhongyan Gong, Ricardo Silva, Alexander N. Flamm, Christopher P. Cook, Lillian Shallow, Jennyffer Souza, Michael Cronce, Christopher J. Arp, Amr A.K. Mousa, Uyen Vo, Cliff Vuong, Jay Keasling, Yalda Moayedi, Jeffery S. Cox, Rithwik Ramachandran, Sarah A. Stanley, Diana M. Bautista

## Abstract

Sneezing, coughing, and pain are common symptoms of respiratory viral infection. These responses are initiated by airway-innervating nociceptors that detect noxious stimuli, but how SARS-CoV-2 activates these neurons remains unclear. Here, we show that the coronavirus papain-like protease (PLpro) directly activates airway-innervating nociceptors in mice and human induced pluripotent stem cell (iPSC)-derived nociceptors. PLpro triggers sneezing, coughing, acute pain, and calcitonin gene-related peptide (CGRP) release in the airways. Both the proteolytic activity of PLpro and the host ion channel transient receptor potential ankyrin 1 (TRPA1) are required for neuronal activation and sneezing. Infection with the mouse-adapted SARS-CoV-2 or the murine coronavirus MHV-A59 also induce sneezing in mice, and MHV-A59-evoked sneezing is TRPA1-dependent. SARS-CoV-2 infection of the trigeminal ganglia altered transcriptional programs associated with nociception, including upregulation of the TRPA1–CGRP signaling pathway activated by PLpro. Together, these findings reveal a direct mechanism by which SARS-CoV-2 triggers pain and airway reflexes and identify PLpro and TRPA1 as potential therapeutic targets for COVID-19.

## Introduction

Common symptoms of viral infections, including SARS-CoV-2, the virus responsible for COVID-19, include sneezing, coughing, headache, and other aches and pain.^1,2^ Sneezing and coughing are host reflexes that may help clear pathogens from the airways but also expel infectious particles that promote transmission to new hosts. Somatosensory and vagal sensory neurons drive these airway reflexes and initiate pain sensations during injury or infection.^3–7^ Yet, to date, no molecular mechanism has been identified by which SARS-CoV-2 or any other virus drives the hallmark symptoms of respiratory infections, such as sneezing and coughing.

When epithelial cells are infected, they release cytokines and other inflammatory mediators that activate surrounding epithelial and immune cells, as well as the sensory neurons that innervate the epithelia. However, infection also generates viral proteins that may be released and modulate host physiology. While viral products are known to activate airway epithelial and immune cells to drive inflammation,^8,9^ little is known about how virus-encoded molecules directly drive respiratory symptoms.

Among viral proteins, proteases have been of much interest as therapeutic targets for COVID-19 due to their critical role in viral replication. SARS-CoV-2 papain-like protease (PLpro) plays an essential role in viral replication by processing viral polyproteins and suppressing the host immune response of infected cells.^10,11^ Proteases from plants, insects, bacteria, and immune cells have been demonstrated to activate sensory neurons to trigger itch, pain, and inflammation,^12–21^ but whether viral proteases do so has been unexplored. PLpro activity can be detected in human throat swabs^22^ and saliva^23^ in individuals with active SARS-CoV-2 infection and at comparable concentrations (29.8 ± 5.9 nM) in supernatants of infected human airway epithelial cells (Figures S1C and S1D). Given that sensory neurons directly innervate the airway epithelial cells that release PLpro during infection, we tested whether PLpro activates airway-innervating sensory neurons. Here, we show that PLpro directly activates sensory neurons via the ion channel TRPA1 to drive sneezing, coughing, and pain.

## RESULTS

### SARS-CoV-2 PLpro activates airway-innervating neurons to drive sneeze, cough, and pain

We first set out to test whether PLpro activates sensory neurons of the nasal mucosa and airways. Using intravital imaging of trigeminal ganglion (TG) sensory neurons, we observed intranasally delivered PLpro-evoked calcium transients (Figure 1A). Instillation of 1 nM PLpro elicited calcium transients in 33.8% of recorded neurons; 21.2% of total neurons were responsive to 1 nM PLpro but not vehicle (Figure 1B). Subsequent instillation of 100 nM and 10 µM PLpro activated an additional 15.3% and 14.3% of neurons, respectively (Figure 1B). Previous physiological studies have shown that stimulation of the nasal mucosa activates both touch-sensitive mechanoreceptors and nociceptors.^24–26^ As expected, instillation of the vehicle or PBS alone activated a subset of airway mechanosensitive neurons (28.4%; Figure S2A); however, these neurons did not exhibit dose-dependent activation in response to PLpro (Figure 1B). These data demonstrate that PLpro uniquely activates a subset of nasal sensory neurons *in vivo* in a dose-dependent manner.

**Figure 1.**
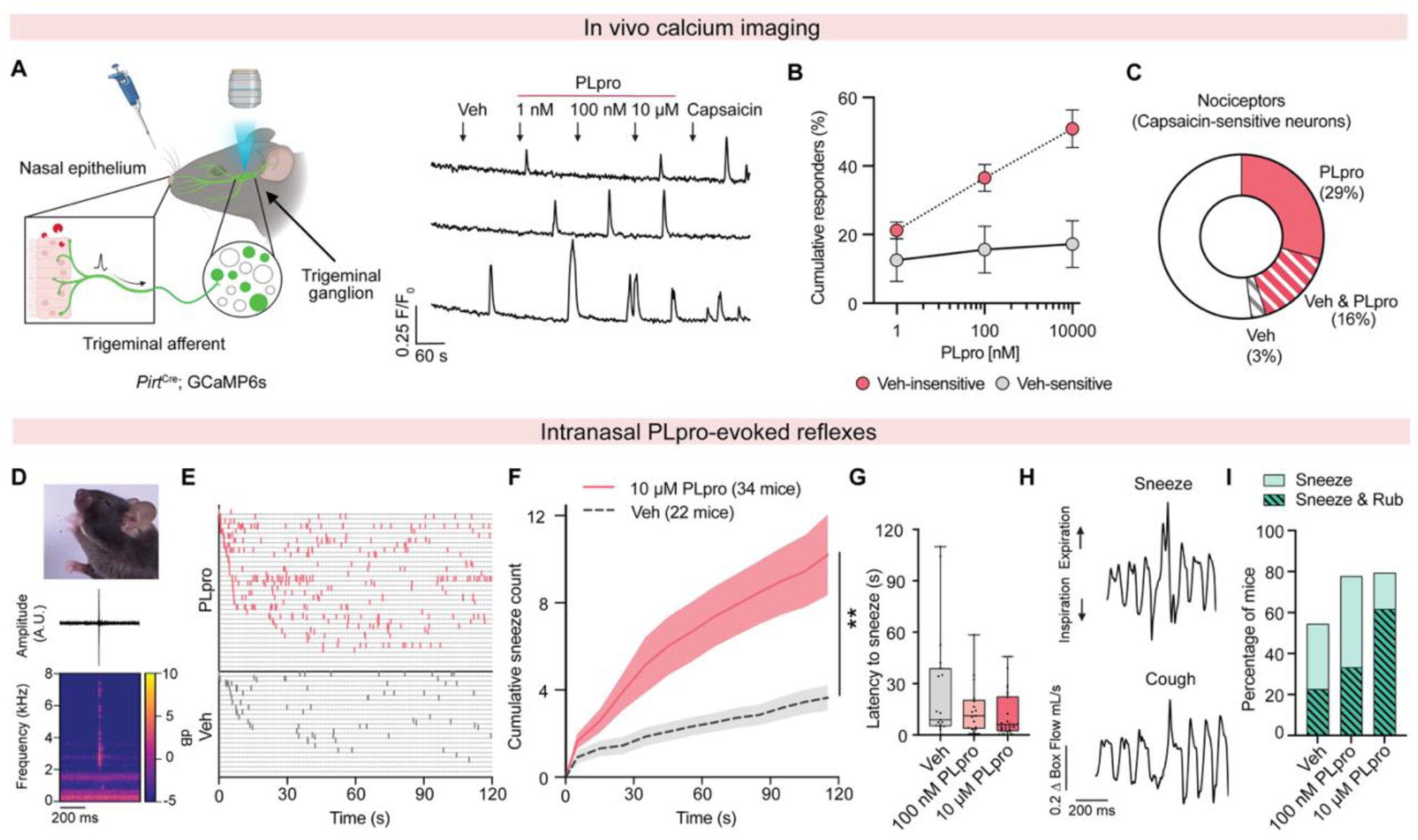
PLpro activates upper airway-innervating neurons to trigger sneeze and cough. (A) Left, schematic of the *in vivo* imaging setup for recording trigeminal ganglion neuron activity during intranasal instillation. Right, representative calcium transients of trigeminal neurons following intranasal instillation of vehicle (Veh), 1 nM, 100 nM, and 10 µM PLpro, and 100 µM capsaicin. Arrowheads indicate time of intranasal instillation. Illustration was created using BioRender (https://www.biorender.com). (B) Percentage of PLpro-responsive neurons among vehicle-sensitive and vehicle-insensitive neurons, *n* = 6 mice. (C) PLpro activates a subset (46/102) of TRPV1+ nociceptors, defined by capsaicin sensitivity; *n=* 6 mice. (D) Intranasal PLpro triggers high-frequency acoustic events (bottom) and nasal expulsions (top). (E) Raster plot of sneeze events, identified by audio recordings, in mice treated intranasally with 10 µM PLpro (*n* = 34 mice) or vehicle (*n* = 22 mice). (F) Mean cumulative sneeze counts following intranasal treatment with 10 µM PLpro or vehicle. Two-way repeated-measures ANOVA: treatment effect, *F*(1, 54) = 7.34, *p* = 0.0090. (G) Latency to the first sneeze following treatment with vehicle (*n* = 22 mice), 100 nM PLpro (*n* = 18 mice), and 10 µM PLpro (*n* = 34 mice). Box plots display the median (center line), 25th–75th percentiles (box), and minimum to maximum values (whiskers). (H) Representative PLpro-evoked sneeze and cough waveforms recorded using whole-body plethysmography. (I) Percentage of mice exhibiting sneezing alone or sneezing accompanied by rubbing within 30 s of treatment. Error bars and shading represent mean ± SEM.

Neurons expressing the ion channel transient receptor potential vanilloid 1 (TRPV1) trigger pain and defensive airway reflexes in response to noxious stimuli, including chemical irritants and inflammatory mediators. Using capsaicin to identify TRPV1+ nociceptors, we found that PLpro activates 45.1% of capsaicin-sensitive neurons (Figure 1C). TRPA1 is expressed in a subset of nociceptors and is a key driver of pain and airway inflammation. Using the TRPA1 agonist allyl isothiocyanate (AITC), we found that 35.6% of AITC-sensitive neurons also respond to PLpro (Figures S2B-S2D). Some PLpro-responsive neurons did not respond to capsaicin or AITC (Figures S2A and S2C). However, sequential stimulus application may cause desensitization that underestimates the overlap between these populations. To assess nociceptor activation in an orthogonal approach, we restricted GCaMP6s expression to Nav1.8+ nociceptors^27^ (*Scn10a*^Cre^). The proportions of PLpro- and AITC-responsive neurons were similar to those observed in the broader *Pirt*^Cre^ sensory neuron population, whereas the proportion of vehicle-responsive neurons was markedly reduced (Figure S2E). These findings indicate that PLpro preferentially activates a subset of nociceptors, although some non-nociceptive neurons may also contribute to the response. By contrast, much of the vehicle response appears to arise from non-nociceptive mechanosensitive neurons. Taken together, these data demonstrate that PLpro acutely activates a subset of airway-innervating nociceptors *in vivo*.

Trigeminal nociceptors mediate sneezing,^28,29^ nasal secretion,^28^ and head pain^30^ common symptoms of viral infection. Which, if any, of these behaviors does PLpro trigger? We used high-speed video and audio recordings to quantify behavioral responses to intranasal treatment of vehicle or PLpro (10 µM). Sneezing and coughing generate distinct sounds in mice and other animals during the expulsive phase, when intrapleural pressure is rapidly released.^3,6,29,31–33^ PLpro elicited high frequency auditory events (Figures 1D and S3A; Videos S1 and S2), typical of a sneeze, over two minutes in mice (Figures 1E and 1F) and shorter latency to sneeze onset (10 µM: 11.6 ± 2.5 s; 100 nM: 15.2 ± 3.7 s) than vehicle-treated mice, which sneezed less and were slower to sneeze (25.9 ± 7.5 s; Figures 1G and S4E-S4H). The sneeze reflex is a forceful expulsion of air that helps clear the airways of irritants and pathogens. To visualize nasal expulsion, we co-applied either PLpro or vehicle combined with blue dye. Indeed, we observed dye expelled onto the sides and floor of the chamber during audible sneeze events (Figures 1D and S3A; Videos S1 and S2). The number of sneezes correlated with the presence of dye at the bottom of the chamber (Figure S3B). By contrast, intranasal administration of 10 µM SARS-CoV-2 main protease (Mpro) did not increase sneezing relative to vehicle treatment (unpaired *t* test: *t* = 1.534, df = 13, *p* = 0.149; Mpro, *n* = 7 mice; vehicle, *n* = 8 mice).

To further characterize the respiratory reflexes triggered by PLpro, we employed whole-body plethysmography and identified specific respiratory patterns that correlated with dye expulsion events (Figure S3C). We observed waveforms stereotypical of sneeze and cough.^3,8,12,37^ Sneeze waveforms consisted of either a single sharp expiratory peak or two in succession, while cough waveforms display an extended inspiration followed by a single expiratory peak (Figure 1H). These findings demonstrate that PLpro triggers rapid sneezing, coughing, and nasal secretion, which could promote expulsion of the virus from the nasal cavity and airways. Previous studies have shown that irritants and inflammatory mediators, such as allergens, histamine, and serotonin, can drive sneezing in mice.^29,34,35^ These data demonstrate for the first time that a viral molecule, PLpro, is alone sufficient to trigger sneezing and coughing, key symptoms of viral infection and putative mechanisms for viral spread.

PLpro induced a dose-dependent increase in sneeze accompanied by a rapid nose rub (Figure S4A). While both 100 nM and 10 µM PLpro rapidly elicited sneezing, nose rubbing was primarily observed with 10 µM PLpro (Figure 1I). In comparison, 100 µM capsaicin, a potent airway irritant, triggered robust sneezing and nose rubbing along with a cessation of face wiping, a response not observed with PLpro (Figures S5A-S5D). Intranasal PLpro did not alter breathing frequency (Figure S4I), suggesting that PLpro triggers acute airway responses in the absence of other respiratory changes. In addition to triggering airway reflexes, activation of trigeminal nociceptors can drive itch or pain behaviors. We thus tested the effects of intradermal injection of PLpro in the cheek, where painful stimuli promote wiping with the forelimbs and itch stimuli evoke scratching with the hindlimb. Mice injected with PLpro displayed rapid wiping behavior, indicative of pain, compared to vehicle-injected mice, but showed no itch-evoked scratching (Figure S6A-S6D). Together, these data demonstrate that PLpro is sufficient to evoke acute nociceptive behaviors associated with irritation and pain in both the airways and skin.

### PLpro directly activates nociceptors via proteolytic activity

To elucidate the mechanisms by which PLpro activates nociceptors, we tested if the protease directly activates sensory neurons *in vitro*. The upper and lower airways are innervated by somatosensory neurons from the trigeminal and dorsal root ganglia (DRG), and vagal sensory neurons from the nodose and jugular ganglia (NJG). We found that PLpro induced calcium transients in neurons cultured from all three ganglia from mice (Figures 2A-2G and Figures S7A-S7C). In DRG cultures, PLpro perfusion elicited rapid, robust calcium transients and a dose-dependent increase in the percentage of neurons reaching a cumulative maximum of 15.2% (Figure 2B). We observed similar proportions of responses in TG cultures (Figure S7A). Interestingly, we observed a greater proportion of neurons that responded to PLpro in cultures of NJG neurons across all PLpro concentrations (33.0% at 100 nM; Figure 2E). Consistent with the activation of nociceptors, PLpro-responsive neurons were of small diameter (Figures 2C, 2F, and S7B) and responded to capsaicin (DRG: 85.7%; NJG: 100%, TG: 63.3%; Figures 2D, 2G, and Figure S7C). A subset of TRPV1+ nociceptors also expresses the TRPA1 channel, and PLpro-responsive neurons in the TG, DRG, and NJG also responded to AITC (DRG: 33.3%, NJG: 50%, TG: 32.9%; Figures 2D, 2G, and Figure S7C). As observed in mouse nociceptors, PLpro and capsaicin also evoked calcium transients in a subset of human induced pluripotent stem cell (iPSC)-derived nociceptors and elicited more calcium influx than vehicle (Figures 2H-2J). Our results show that PLpro directly activates a subset of human and mouse nociceptors.

**Figure 2.**
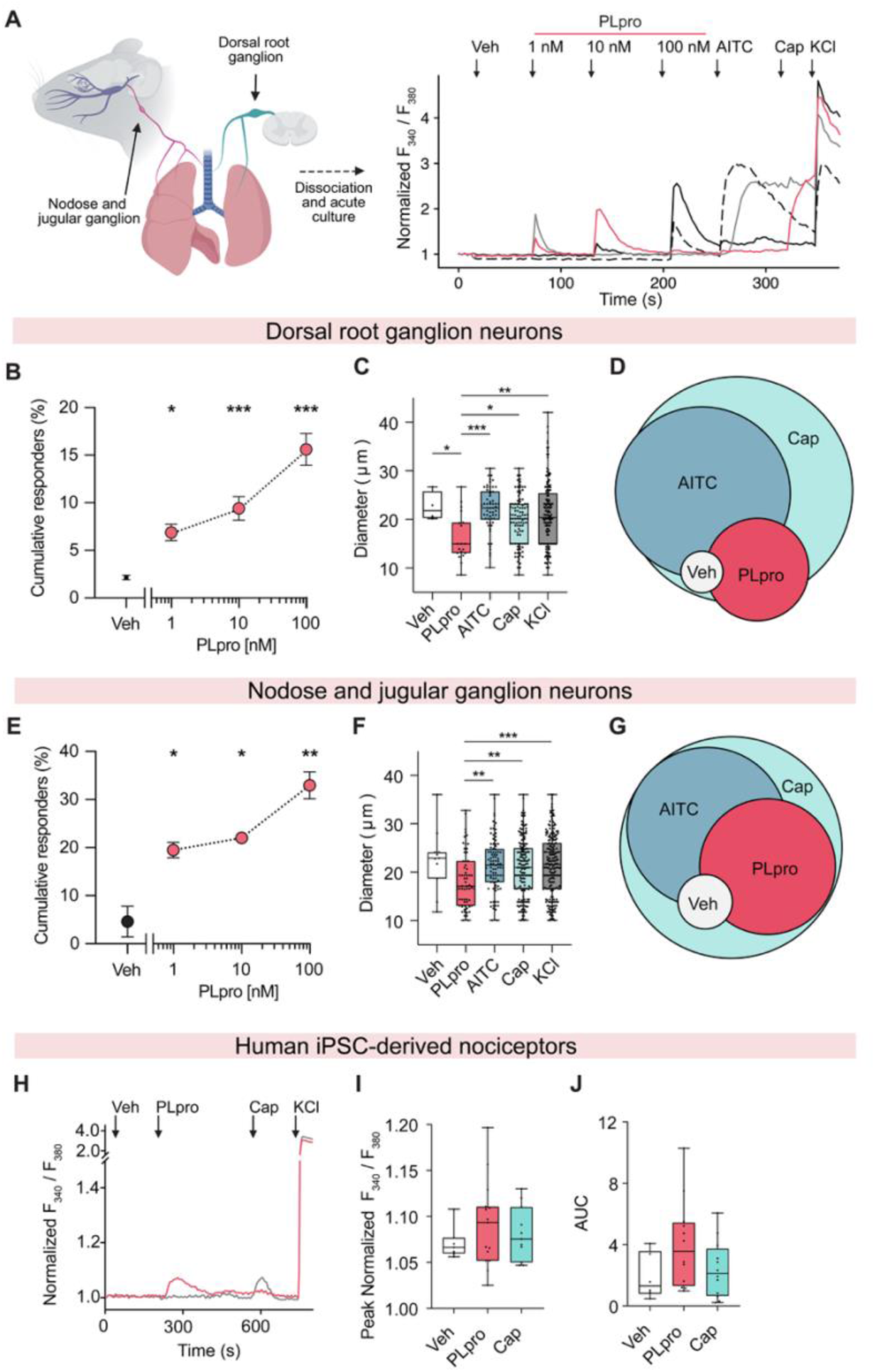
PLpro directly activates nociceptors. (A) Left, schematic illustrating airway-innervating sensory ganglia. Right, representative calcium transients from cultured dorsal root ganglion (DRG) neurons following perfusion of vehicle (Veh), 1 nM, 10 nM, 100 nM PLpro, 100 µM allyl isothiocyanate (AITC), 1 µM capsaicin (Cap), and high-potassium solution (KCl). Arrowheads indicate time of stimulus perfusion. Illustration was created using BioRender (https://www.biorender.com). (B) PLpro induces dose-dependent activation of DRG neurons: 1 nM (6.9%), 10 nM (9.4%), and 100 nM (15.6%), significantly greater than vehicle (2.2%). One-way ANOVA: *F*(3, 36) = 24.0, *p* < 0.0001; Dunnett’s post hoc test: *p* (Veh vs. 1 nM) = 0.0168, *p* (Veh vs. 10 nM) = 0.0002, *p* (Veh vs. 100 nM) < 0.0001, *n* = 10 mice. (C) Diameters of stimulus-responsive DRG neurons. PLpro-responsive (combined responders across 1, 10, and 100 nM) neurons were smaller than Veh-, AITC-(100 µM), Cap-(1 µM), and KCl-responsive neurons; Welch’s ANOVA: *F*(4, 21.42) = 6.11, *p* = 0.0019; Dunnett’s T3 test vs. PLpro: Veh, *p* = 0.0390; AITC, *p* = 0.0003; Cap, *p* = 0.0268; KCl, *p* = 0.0027. (D) Venn diagram of PLpro-, Veh-, AITC-, and Cap-responsive DRG neurons (*n* = 154 neurons). (E) PLpro induces dose-dependent activation of cultured nodose and jugular ganglion (NJG) neurons: 1 nM (19.6%), 10 nM (22.0%), and 100 nM (33.0%), significantly greater than vehicle (4.6%). One-way ANOVA: *F*(3, 4) = 25.50, *p* = 0.0046; Dunnett’s post hoc test: *p* (Veh vs. 1 nM) = 0.0235, *p* (Veh vs. 10 nM) = 0.0134, *p* (Veh vs. 100 nM) = 0.0022, *n* = 2 biological replicates, each containing NJG from 2 mice. (F) Diameters of stimulus-responsive NJG neurons. PLpro-responsive neurons were smaller than AITC-, Cap-, and KCl-responsive neurons; Welch’s ANOVA: *F*(4, 66.18) = 4.20, *p* = 0.0043; Dunnett’s T3 test vs. PLpro: Veh, *p* = 0.2044; AITC, *p* = 0.0024; Cap, *p* = 0.0084; KCl, *p* = 0.0007. (G) Venn diagram of PLpro-, Veh-, AITC-, and Cap-responsive NJG neurons (*n* = 220 neurons). (H) Representative calcium transients from human iPSC-derived nociceptors during perfusion of vehicle, 500 nM PLpro, 1 µM capsaicin, and KCl. (I and J) (I) Peak normalized calcium responses and (J) area under the curve (AUC) following perfusion of vehicle, PLpro, or capsaicin (*n* = 4 wells per condition from two independent differentiations). Biological replicates (individual mouse or pooled mice) contain 3–4 wells containing ∼100–300 neurons per well. Error bars represent mean ± SEM. Box plots display the median (center line), 25th–75th percentiles (box), and minimum to maximum values (whiskers).

Proteolytic activity of PLpro is required for both viral protein processing and inhibition of anti-viral immunity.^11^ Is proteolytic activity also required for neuronal activation? Studies have shown that a point mutation of the key catalytic cysteine to a serine (PLpro_C111S_) results in a catalytically inactive mutant PLpro (Figures 3A and 3B). The PLpro_C111S_ mutant activated significantly fewer DRG neurons than active PLpro_WT_ *in vitro* (Figure 3D), comparable to activation by vehicle perfusion. Likewise, co-application of PLpro with the PLpro inhibitor 19 that blocks catalytic activity^36^ significantly decreased the percentage of DRG neurons that are activated (Figure S7E). Together, this demonstrates that neuronal activation by PLpro is dependent on its cysteine protease activity. Thus, we asked whether catalytic activity is also required for PLpro-evoked sneezing. Indeed, unlike treatment with PLpro, intranasal treatment with the PLpro_C111S_ mutant elicited few sneezes, similar to vehicle treatment (Figures 3E and 3F). These data show that both neuronal activation and sneeze reflexes are mediated directly by the proteolytic activity of PLpro.

**Figure 3.**
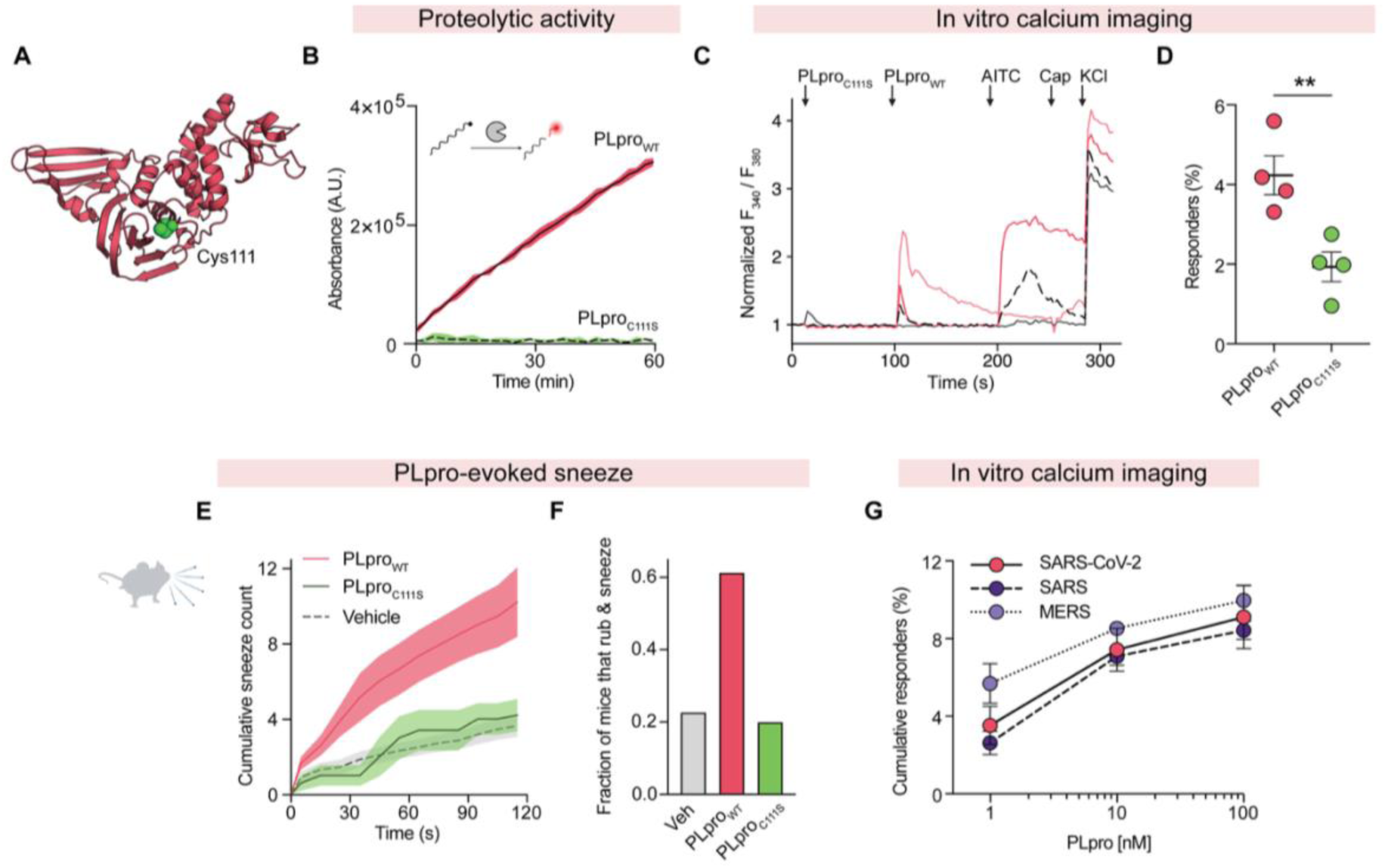
PLpro catalytic activity is required for sensory neuron activation and sneezing. (A) Structure of PLpro (PDB: 7JIW, pdb_00007jiw) highlighting the active site residue Cys111 (represented as spheres). (B) Proteolytic activity measured by fluorescence from cleaved substrate for WT SARS-CoV-2 PLpro (PLpro_WT_) and catalytically inactive mutant PLpro (PLpro_C111S_). (C) Representative calcium transients from cultured DRG neurons following perfusion of 100 nM PLpro_C111S_, 100 nM PLpro_WT_, 100 µM AITC, 1 µM capsaicin, and high potassium solution (KCl). Arrowheads indicate time of perfusion. (D) PLpro_C111S_ activates significantly fewer neurons compared to PLpro_WT_ (unpaired *t*-test: *t* = 3.755, *df* = 6, *p* = 0.0095, *n* = 4 mice). (E) Average cumulative sneeze counts following intranasal treatment of vehicle (*n* = 22 mice), 10 µM PLpro_WT_ (*n* = 34 mice), and 10 µM PLpro_C111S_ (*n* = 5 mice). (F) Fraction of mice exhibiting rubbing and sneezing behaviors within 30 seconds of intranasal treatment of vehicle, 10 µM PLpro_WT_, and 10 µM PLpro_C111S_. (G) PLpro orthologs from SARS-CoV-2, SARS-CoV (SARS), and MERS-CoV (MERS) induce nearly identical dose-dependent activation of cultured DRG neurons. Two-way ANOVA: *F*(1.853, 11.12) = 90.91, *p* (dose) < 0.0001, *F*(2,6) = 0.7545, *p* (coronavirus species) = 0.5455, *F*(4,12) = 0.8069, *p* (interaction) = 0.5101, SARS-CoV-2 (*n =* 4 mice), SARS (*n =* 3 mice), and MERS (*n =* 2 mice), with 3–4 wells per animal containing ∼100– 300 neurons per well. Error bars and shading represent mean ± SEM. Illustration in E was created using BioRender (https://www.biorender.com).

Papain-like proteases are expressed by all coronaviruses, including across diverse species and other human pathogenic betacoronaviruses. SARS-CoV (SARS) and MERS-CoV (MERS) share 89% and 51% sequence similarity with SARS-CoV-2 PLpro, respectively, and these proteases each display different substrate specificities and catalytic efficiencies.^11,36,37^ Despite these differences, all three proteases recognize a conserved L-x-G-G cleavage motif. We therefore asked whether these divergent PLpro orthologs could also activate mouse neurons. SARS, MERS, and SARS-CoV-2 PLpro elicited nearly identical dose-dependent activation of cultured DRG neurons (Figure 3G), suggesting that their conserved neuronal activity may reflect shared features of PLpro substrate recognition, including recognition of the conserved cleavage motif. This L-x-G-G cleavage motif is also conserved in the murine hepatitis virus (MHV) papain-like protease 2 (PLP2), a betacoronavirus that naturally infects mice.^38,39^ Together, these data demonstrate that direct neuronal activation via PLpro is evolutionarily conserved among pathogenic betacoronaviruses.

### The host TRPA1 ion channel mediates PLpro- and coronavirus-evoked sneezing

PLpro opens a calcium-permeable ion channel in the plasma membrane, as calcium responses were eliminated when extracellular calcium was chelated (Figure S7D). Airway irritants and inflammatory mediators, including proteases, activate nociceptors via the calcium-permeable ion channels, TRPV1 or TRPA1.^40–44^ To test the requirement for these channels, we examined PLpro responses in cultured DRG neurons from TRPV1- or TRPA1-deficient mice and found that the percent of PLpro responders was reduced (wild type [WT]: 6.8%, *Trpv1* knockout [KO]: 4.3%, *Trpa1* KO: 2.7%; Figures 4A and S7D). The acute blockade of TRPA1 with the antagonist HC-030031 (HC) but not blockade of TRPV1 with the antagonist AMG 517 (AMG) also reduced PLpro responders *in vitro* (control: 4.4%, AMG: 7.4%, HC: 2.5%; Figures 4B and S7D), comparable to responses observed in *Trpa1* KO neurons. These data demonstrate that PLpro activates sensory neurons via the TRPA1 ion channel.

**Figure 4.**
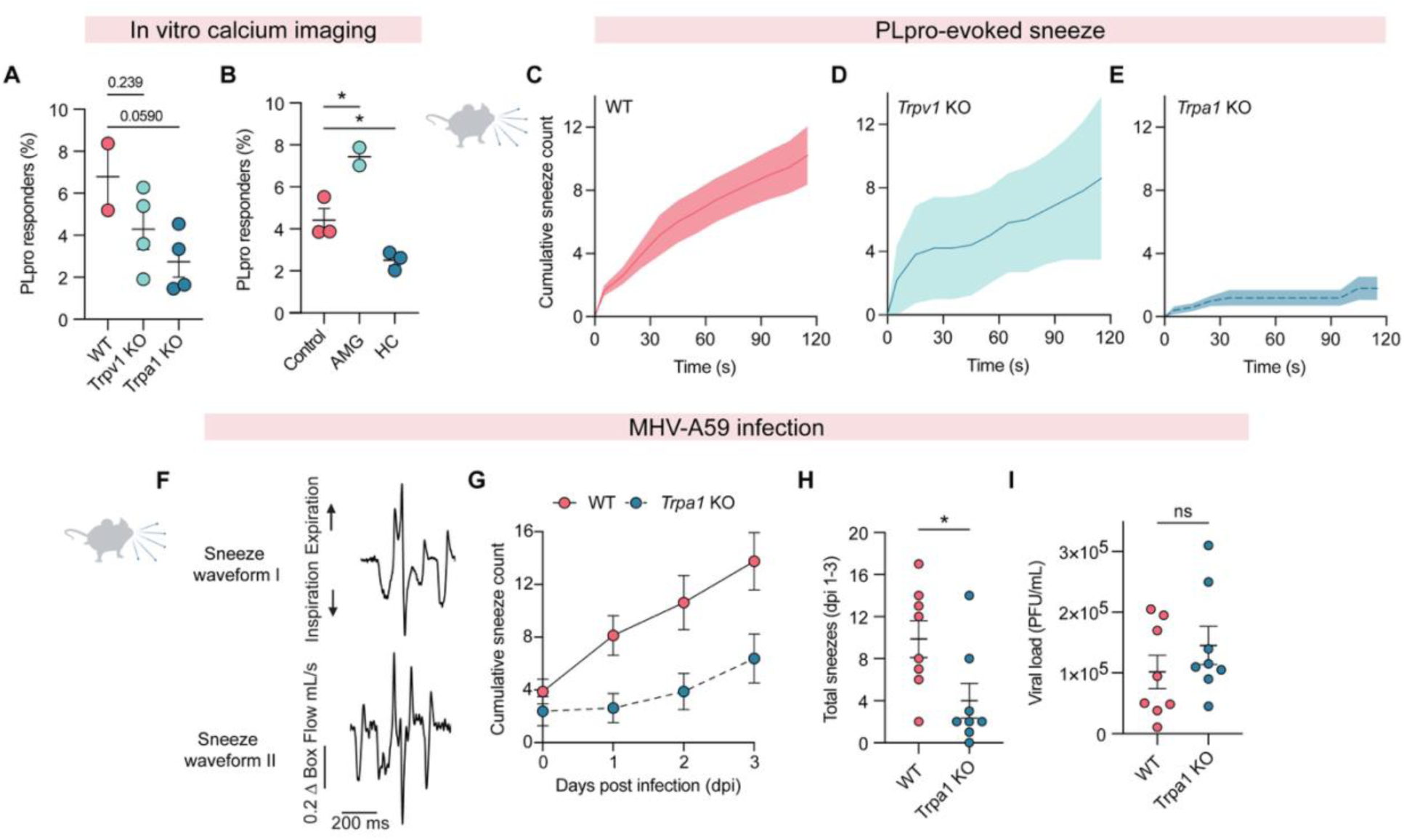
PLpro and coronavirus-evoked sneezing require TRPA1. (A) Percentage of cultured DRG neurons responding to 100 nM PLpro *in vitro* from wild-type C57BL/6J (WT) mice (*n =* 2 mice), *Trpv1* KO mice (*n =* 4 mice), and *Trpa1* KO mice (*n =* 4 mice). One-way ANOVA: *F*(2, 7) = 3.367, *p* = 0.0945; Dunnett’s post hoc test: *p* (WT vs. *Trpv1* KO) = 0.2386, *p* (WT vs. *Trpa1* KO) = 0.0590. (B) Percentage of DRG neurons from WT mice responding to 100 nM PLpro in the presence of vehicle (control; *n =* 3 mice), a TRPV1 antagonist AMG-517 (AMG; *n =* 2 mice), or a TRPA1 antagonist HC-030031 (HC; *n =* 3 mice). One-way ANOVA: *F*(2, 5) = 28.48, *p* = 0.0019; Dunnett’s post hoc test: *p* (Veh vs. AMG) = 0.0102, *p* (Veh vs. HC) = 0.0389. (C–E) Cumulative sneeze counts following intranasal treatment of 10 μM PLpro in (C) WT (*n =* 34 mice), (D) *Trpv1* KO mice (*n =* 5 mice), and (E) *Trpa1* KO mice (*n =* 5 mice). (F) Representative whole-body plethysmography traces of two sneeze waveforms observed following MHV-A59 infection. (G) Cumulative sneeze counts in WT and *Trpa1* KO mice from 0 to 3 days post-infection (dpi). (H) Total sneezes from 1 to 3 dpi were reduced in *Trpa1* KO mice compared with WT mice, unpaired *t* test, *t* = 2.439, *df* = 14, *p* = 0.0286, *n* = 8 mice per genotype. (I) Lung viral titers at 3 dpi did not differ between WT and *Trpa1* KO, unpaired *t* test, *t* = 1.057, *df* = 14, *p* = 0.3082, *n* = 8 mice per genotype. Data in (G–I) are from the same cohort; Data are mean ± SEM.

We thus asked whether PLpro directly activates TRPA1. Unlike AITC, PLpro failed to evoke currents in HEK293T cells heterologously expressing human TRPA1 (Figures S8A-S8C). This is consistent with the absence of the canonical PLpro cleavage motif in both human and mouse TRPA1 and suggests that PLpro engages TRPA1 via an intermediate receptor or signaling pathway. Protease-activated receptors (PARs) and Mas-related G protein-coupled receptors (MRGPRs) are candidate upstream targets; however, like TRPA1, these too lack the canonical PLpro cleavage motif, and heterologously expressed receptors are not activated by PLpro (Figures S8D-S8J). Thus, although TRPA1 is required for PLpro-induced neuronal activation, the receptor and signaling pathway coupling PLpro activity to TRPA1 remain to be determined.

Given that TRPA1 is required for neuronal activation *in vitro*, we assessed the requirement of TRPA1 for PLpro-evoked sneezing. Intranasal PLpro triggers robust sneezing in WT and TRPV1-deficient mice but not in TRPA1-deficient mice (Figures 4C-4E). However, TRPA1-deficient mice are still capable of sneezing, as they display sneezing responses to capsaicin comparable to those of WT mice, indicating that the sneeze reflex itself is intact (Figure S5E). We next examined whether coronavirus infection similarly elicits sneezing in mice. Mice infected intranasally with either the mouse coronavirus murine hepatitis virus strain A59 (MHV-A59) or mouse-adapted MA10 SARS-CoV-2 exhibited sneeze and cough responses over the course of infection consistent with infection of both the upper and lower airways at this volume (40 µL; Figure S9A-S9H). We therefore asked whether infection-evoked sneezing, like PLpro-evoked sneezing, required TRPA1. Following intranasal infection with MHV-A59, TRPA1-deficient mice exhibited significantly fewer sneezes than WT mice (Figures 4G and 4H). Importantly, the loss of sneeze occurred despite comparable viral loads, mortality, weight loss, and respiratory depression (Figure 4I and Figures S9I-S9K). Thus, TRPA1 is required for driving sneeze responses during coronavirus infection but is dispensable for viral control, disease severity, and sickness behaviors 3 days post infection. Although TRPA1 is well known to mediate cough and inflammatory pain,^40,45^ our findings identify a role for TRPA1 in sensing a viral protease and driving sneezing, a key host response to respiratory infection.

### PLpro and SARS-CoV-2 infection converge on a TRPA1-CGRP signaling pathway

In addition to sneezing, pain is a common symptom of acute COVID-19 infection, and animal models of SARS-CoV-2 infection display mechanical allodynia both during and after infection.^1,46,47^ A wide variety of irritants and inflammatory mediators induce mechanical or thermal sensitization via TRPV1 and TRPA1.^40,48^ Given that PLpro acutely activates nociceptors and evokes nocifensive behaviors (Figures S6A and S6B), we tested if PLpro could also drive pain hypersensitivity. We found that PLpro promotes mechanical hypersensitivity following intradermal injection into the hindpaw. PLpro significantly reduced the von Frey paw withdrawal threshold 3 h after injection that returned to baseline by 24 h (Figure 5A). TRPA1-deficient mice failed to develop the PLpro-induced mechanical sensitization observed in WT and TRPV1-deficient mice (Figure 5B and Figures S6E-S6G). Interestingly, we observed no difference in heat sensitivity between vehicle- and PLpro-injected paws (Figure 5C), consistent with TRPA1’s role in promoting mechanical but not thermal sensitization.^40,49^

**Figure 5.**
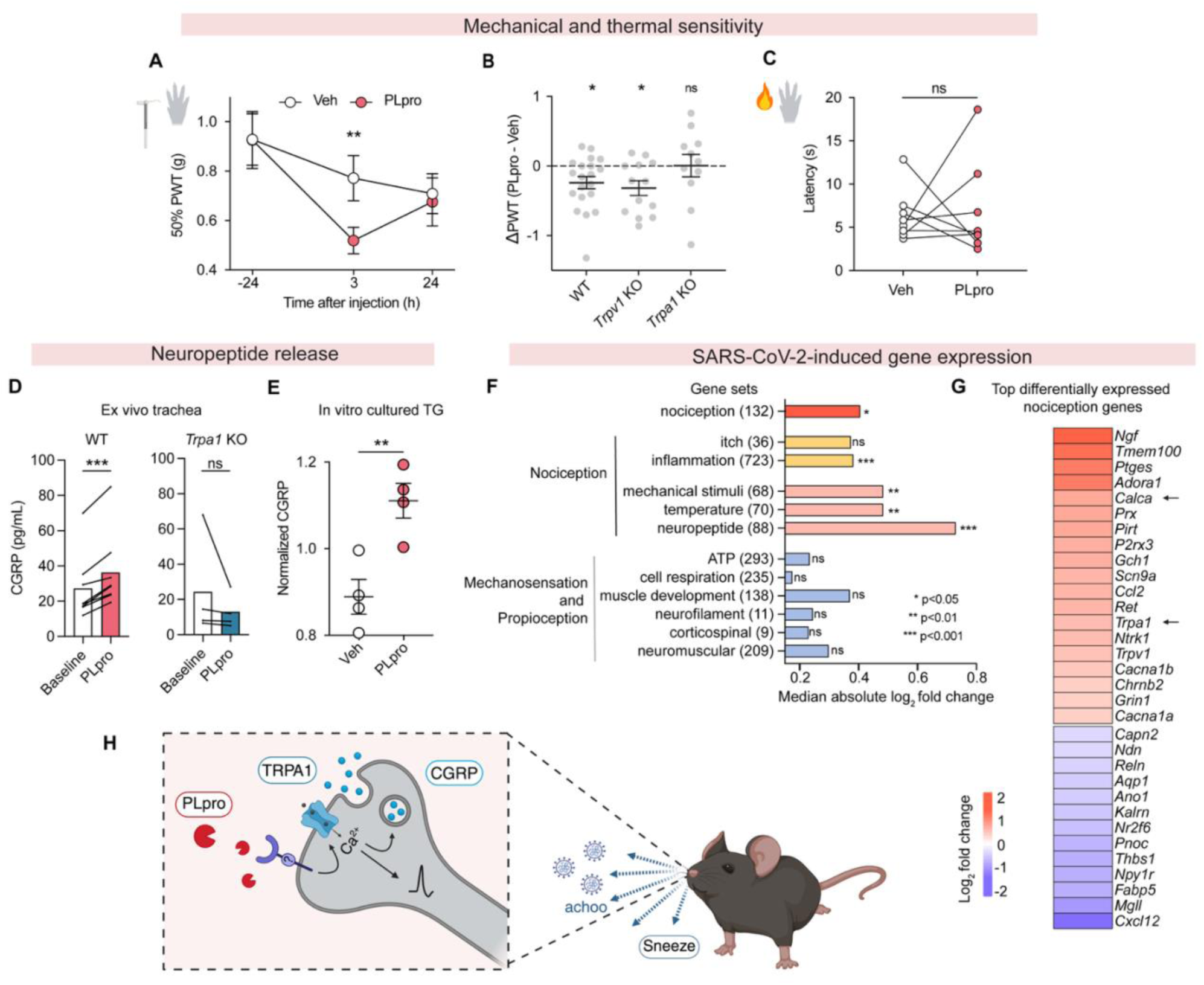
SARS-CoV-2 infection and PLpro activation share a key nociceptive signature. (A) Intradermal injection of 50 μM PLpro reduces the paw withdrawal threshold (PWT) at 3 h post-injection but not at 24 h, compared with baseline (24 h pre-injection). Two-way repeated-measures ANOVA: *F*(1.793, 64.56) = 5.477, *p* (time) = 0.008; Dunnett’s multiple comparisons: Vehicle-injected paws: *p* (Baseline vs. 3 h) = 0.5550, *p* (Baseline vs. 24 h) = 0.0729; PLpro-injected paws: *p* (Baseline vs. 3 h) = 0.0077, *p* (Baseline vs. 24 h) = 0.2007; *n* = 19 mice per group. (B) WT and *Trpv1* KO mice, but not *Trpa1* KO mice, develop mechanical hypersensitivity 3 h post-injection. WT: one-sample *t* test, *t* = 2.746, *df* = 19, *p* = 0.013, *n* = 20 mice; *Trpv1* KO: *t* = 3.001, *df* = 12, *p* = 0.011, *n* = 13 mice; *Trpa1* KO: *t* = 0.039, *df* = 10, *p* = 0.970, *n* = 11 mice. (C) Intradermal PLpro injection does not alter response latency to radiant heat (paired *t* test: *t* = 0.233, *df* = 7, *p* = 0.822, *n* = 8 mice). (D) PLpro significantly increases CGRP release compared with baseline in an *ex vivo* tracheal preparation in WT but not *Trpa1* KO mice. WT: paired *t* test: *t* = 6.878, *df* = 7, *p* = 0.0002, *n* = 8 mice. *Trpa1* KO mice: *t* = 1.158, *df* = 3, *p* = 0.3308, *n* = 4 mice. (E) 100 nM PLpro induces CGRP release from cultured TG: unpaired *t* test: *t* = 3.928, *df* = 6, *p* = 0.008, *n* = 4 mice. (F) SARS-CoV-2 infection significantly alters gene sets associated with nociception. Median absolute log_2_ fold changes for each custom gene set were tested against 100,000 random permutations of identically sized gene sets (gene set sizes listed in parenthesis). Mock-infected: *n* = 4 wells, SARS-CoV-2-infected: *n* = 3 wells. *p < 0.05, **p < 0.01, ***p < 0.001. (G) Heatmap of the most significantly differentially expressed nociception-associated transcripts (log_2_ fold change: infection vs. mock). (H) Model: PLpro activates nociceptors through its proteolytic activity, triggering TRPA1-dependent sneezing, coughing, and pain behaviors. Illustration was created using BioRender (https://www.biorender.com). Data are mean ± SEM.

A key hallmark of TRPA1 activation is the release of the neuropeptides calcitonin gene-related peptide (CGRP) and Substance P that can drive neurogenic inflammation^50^ and immunomodulation.^4,51^ We therefore asked whether PLpro was sufficient to elicit neuropeptide release in the airways. Using an *ex vivo* mouse tracheal preparation, we found that PLpro induced significant CGRP release above baseline in WT mice (1.4-fold relative to baseline; Figure 5D). This release occurred in a sensory neuron-dependent manner, as both global *Trpa1* KO mice and mice lacking *Trpa1* specifically in sensory neurons (*Pirt*^Cre^ *;Trpa1*^fl/fl^) displayed no significant CGRP release (Figures 5D and S10A). Consistent with these findings, PLpro also triggered acute CGRP release from cultured trigeminal ganglia cells *in vitro* (Figure 5E). Surprisingly, unlike allergen proteases, PLpro did not induce detectable Substance P release (Figure S10B).^17,18^ Together these data demonstrate that PLpro activates TRPA1 signaling in sensory neurons to selectively drive acute CGRP release both *in vitro* and in the airways. CGRP has been detected in bronchoalveolar lavage fluid of SARS-CoV-2 infected patients and correlates with viral load and disease severity.^52^ Our findings provide a potential mechanism linking SARS-CoV-2 PLpro activity to CGRP release in the respiratory tract.

Moreover, SARS-CoV-2 infection has been observed in the sensory ganglia of humans, hamsters, and mice.^46,53–56^ While sensory neurons are susceptible to infection, unlike in epithelial cells, the virus does not replicate.^53–55^ Despite this lack of replication, infection drives gene expression changes in hamster sensory ganglia that are related to neuronal signaling and neuroinflammation,^46^ which could reflect consequences of the systemic response to infection or direct infection of the ganglia. Thus, we assessed the consequences of direct SARS-CoV-2 infection on cultured mouse trigeminal ganglia. SARS-CoV-2 infection significantly upregulated 789 genes and downregulated 1664 genes, with the most differentially expressed genes being related to inflammation (Figures S10C and S10D). At the whole-ganglion level, nociception-associated gene sets were significantly altered, including transcriptional signatures associated with both non-peptidergic and peptidergic nociceptors but not those associated with mechanosensation, proprioception, and itch (Figure 5F).^57^ Because trigeminal ganglia cultures contain neurons, glia, and resident immune cells, these bulk transcriptional changes cannot be attributed to a specific cell type. Within this bulk transcriptional response, among the most upregulated genes were several ion-channel genes (*P2rx3, Scn9a, Trpa1, Trpv1, Cacna1b, Cacna1a, Chrnb2,* and *Grin1*) associated with nociceptor excitability and pain signaling (Figure 5G). SARS-CoV-2 infection also upregulated *Calca* and *Calcb,* the genes that encode CGRP-ɑ and -β, but not *Tac1,* the gene that encodes Substance P (Figure S10F). The same TRPA1-CGRP signaling pathway that is directly activated by PLpro is also transcriptionally upregulated by SARS-CoV-2 infection. We found that treatment with PLpro alone did not induce any significantly differentially expressed genes (Figure S10G), suggesting that infection-associated transcriptional changes arise through a mechanism distinct from acute activation by PLpro. Together, these findings reveal two complementary mechanisms by which SARS-CoV-2 targets nociceptor signaling: PLpro-dependent activation and infection-induced changes in nociception-related transcriptional programs.

## DISCUSSION

Symptoms of viral infection, such as sneezing, coughing, and headache, result from activation of somatosensory and vagal sensory neurons. These symptoms have largely been attributed to the actions of inflammatory mediators released by infected epithelial and immune cells on neurons. Here, we show that SARS-CoV-2 can directly activate sensory neurons through PLpro. This activity is conserved across PLpro orthologs from SARS-CoV and MERS-CoV. By demonstrating that PLpro directly activates sensory neurons to trigger sneezing and coughing, independent of inflammation, our study expands our understanding of how pathogens engage the nervous system to shape host physiology. Together with recent studies showing that *Mycobacterium tuberculosis* directly activates neurons to trigger cough,^58^ *Staphylococcus aureus* targets neurons to suppress lung immunity,^59^ bacterial products like lipopolysaccharide engage airway neurons to drive sickness behaviors,^60^ and SARS-CoV-2 spike protein activates vagal neurons,^61^ our findings suggest that nociceptors are likely widespread yet underappreciated targets of respiratory pathogens.

We find that PLpro activates sensory neurons via the calcium-permeable ion channel TRPA1. This neuronal activation requires proteolytic activity of PLpro, which is also required for polyprotein processing during viral replication and for suppression of host antiviral responses through ISG15 cleavage. Our findings therefore identify a third function for PLpro: directly activating sensory neurons to trigger sneezing and coughing. How PLpro proteolytic activity is coupled to TRPA1 activation remains unclear. We show that PLpro did not directly activate heterologously expressed TRPA1 or known protease-activated receptors (PARs and MRGPRs), suggesting the involvement of an unidentified intermediate receptor or signaling mechanism. Identifying the direct target of PLpro will be an important direction for future work and may reveal a new pathway by which viral proteases, and potentially proteases from a diversity of organisms, engage sensory neurons.

Importantly, our functional studies show that TRPA1 is required for sneezing evoked by both PLpro and coronavirus infection, establishing TRPA1 as a critical transducer of host sensory responses to coronaviruses. TRPA1 is a key sensor for irritants and drives protective reflexes that help clear harmful stimuli from the airways. The requirement for TRPA1 in MHV-A59 infection-triggered sneezing, together with the conserved neuronal activation of PLpro orthologs, raises the possibility that the host may have evolved to recognize a conserved feature of coronavirus infection to trigger an airway defense response. Conversely, because sneezing and coughing can aerosolize and disperse viral particles, coronaviruses may also benefit from engaging this pathway. This raises a key question as to whether sneezing and coughing during infection is a host defense mechanism that clears pathogens from the airways or whether these reflexes are exploited by the virus as a means of transmission. Future studies dissecting the contribution of these reflexes to pathogen clearance and viral transmission will be important for resolving this question. Thus, PLpro represents an attractive therapeutic target for COVID-19, as PLpro inhibitors may both suppress viral replication while also alleviating virus-induced symptoms such as sneezing, coughing, and pain.

## MATERIALS AND METHODS

### Experimental model and study participant details

#### Mice

All mice were housed in standard conditions in accordance with standards approved by the Animal Care and Use Committee of the University of California Berkeley (14 hour light: 10 hour dark cycle, 21 °C). Wild-type C57BL/6N mice were obtained from Charles River for neonatal trigeminal cultures and C57BL/6J from Jackson Laboratories and were used for all other mouse experiments. All experiments were performed under the policies and recommendations of the International Association for the Study of Pain and approved by the University of California Berkeley Animal Care and Use Committee. Where appropriate, genotyping was performed by Transnetyx using real-time PCR. Mouse lines used in this study included *Pirt*^Cre^ (Pirttm3.1(cre)Xzd) gift from Dr Xinzhong Dong (Johns Hopkins University, Baltimore, MD) and Ai96 (B6J.Cg-Gt(ROSA)26Sortm96^(CAG-GCaMP6s)Hze^/MwarJ; Jackson Laboratories 028866), *Trpv1* KO (Trpv1tm1Jul RRID:MGI:4417977; Jackson Laboratories 003770), and *Trpa1* flox (B6.129S-Trpa1tm2Kykw/J; Jackson Laboratories 008650), *Scn10a*^Cre^ (B6.129(Cg)-Scn10atm2(cre)Jwo/TjpJ; Jackson Laboratories 036564), and *Trpa1* KO (Trpa1tm1Jul, RRID:MGI:3696956) originally from Jackson Laboratories were bred in-house.

For *in vitro* cultures of sensory ganglia (TG, DRG, and NJG), male and female mice aged 6-16 weeks were used, and data from both sexes were combined. Neonatal TG cultures were prepared from mice postnatal days 1-5. *In vivo* calcium imaging experiments were performed in male and female mice aged 8-40 weeks. For behavioral experiments, male and female mice aged 8-12 weeks were used. No significant difference in acute sneeze behavior was observed between male and female mice (Figure S11D); therefore, data from both sexes were combined. Infection experiments were performed in male mice aged 8-10 weeks; therefore, whether sex influences infection-induced sneezing remains to be determined.

#### Primary cell culture

Trigeminal ganglia (TG), dorsal root ganglia (DRG) and nodose and jugular ganglia (NJG) were cultured as previously described.^42,62^ In brief, sensory ganglia were dissected and incubated in warmed Collagenase P (Roche) in Hanks calcium-free balanced salt solution (HBSS) followed by incubation in 0.25% trypsin with gentle agitation. Cells were washed, triturated, and plated in Minimum Essential Medium (MEM, Gibco) supplemented with 10% horse serum (v/v), MEM vitamins, penicillin, streptomycin and L-glutamine. Neonatal TG cultures were plated on poly-D-lysine coated eight-well chambers for calcium imaging. Adult TG, DRG, and NJG were plated on poly-D-lysine and laminin-coated eight-well chambers for calcium imaging or 96-well plates for infection and ELISA experiments. Age-matched WT controls were prepared on the same day for experiments on neurons from knockout mice.

#### Human induced pluripotent stem cells

Human iPSC-derived neurons were differentiated per manufacturer’s instructions and used in calcium imaging 21 and 39 days after differentiation (Axol Bioscience, ax0555). Neuronal viability and functionality were validated by responses to depolarization evoked by high potassium solution (75 mM KCl) and further confirmed by capsaicin-evoked calcium transients and depolarization-evoked substance P release.

#### Cell culture

Human adenocarcinoma lung epithelial (Calu-3) cells (ATCC, HTB-55) were maintained with RPMI (Fisher Scientific) supplemented with fetal bovine serum (FBS), L-glutamine, penicillin/streptomycin and maintained in a humidified incubator at 37 °C with 5% CO_2_. HEK293T cells (CRL-3216, ATCC) were cultured in complete growth medium consisting of DMEM (11960044, Gibco) supplemented with 2 mM L-glutamine (25030081, Gibco) and 10% FBS (SH30396.03, HyClone) and maintained in a humidified incubator at 37 °C with 5% CO_2_.

### Method Details

#### *In vivo* calcium imaging of trigeminal ganglion

*In vivo* calcium imaging of trigeminal neurons was conducted in anesthetized mice as previously described.^63^ In brief, *Pirt*^Cre^;GCaMP6s mice were anesthetized with ketamine and xylazine (100 and 10 mg/kg intraperitoneal) and body temperature was maintained at 36°C throughout surgery and imaging. Mice were head-fixed on a custom chamber and tracheotomized to permit breathing while reducing aspiration into the lower airways. A unilateral craniotomy and hemispherectomy was performed to expose the dorsal surface of the right trigeminal ganglion. Hemostasis was achieved using Surgifoam gel sponge (Ethicon 1972). Images were collected at 5 Hz using a Sutter Movable Objective Microscope (MOM) equipped with an Olympus XL-Fluor 4X/340 (NA 0.28) lens and ORCA-Fusion BT Digital CMOS camera. 480 nm illumination was provided by Lambda 721 (Sutter). Stimuli were applied via intranasal instillation of 10 µL of solution applied evenly between both nostrils using a P20 pipette. Stimuli were applied at 2-minute intervals.

#### *In vitro* calcium imaging

Neuronal calcium imaging experiments were carried out as previously described.^62^ Neuronal cultures were loaded for 45 minutes at room temperature with 10 µM Fura-2AM supplemented with 0.01% pluronic F-127 (wt/vol, Life Technologies) in a physiological Ringer’s solution containing (in mM) 140 NaCl, 5 KCl, 10 HEPES, 2 CaCl_2_, 2 MgCl_2_ and 10 D-(+)-glucose, pH 7.4. Neurons were stimulated with SARS-CoV-2 PLpro (R&D #E-611), SARS-CoV-2 PLpro C111S mutant (R&D custom order), SARS-CoV PLpro (R&D #E-610), MERS-CoV PLpro (R&D #E-609), or SARS-CoV-2 PLpro (Cayman #31817) diluted in Ringer’s to the appropriate concentration. Supplier, preparation method, and purity for each protease used in this study are provided in Table S1. The vehicle (50 mM HEPES, 300 mM NaCl, 1 mM TCEP, 10% (v/v) glycerol, pH 8.0) was matched to the dilution buffer for PLpro (R&D #E-611). For PLpro (Cayman #31817), Ringer’s solution served as the vehicle. No differences in neuronal activation were observed between the two PLpro preparations or their respective vehicles, so data were combined (Figure S11C). Cells were identified as neurons by eliciting depolarization with high potassium Ringer’s solution (75 mM) at the end of each experiment. Acquired images were displayed as the ratio of 340 nm / 380 nm. Fura-2 ratio was normalized to baseline ratio.

#### MRGPRX1 and MRGPRX2 calcium imaging

Chem-1 cells stably expressing MRGPRX1 (Eurofins Discoverx HTS122RTA) and MRGPRX2 (Eurofins Discoverx HTS058RTA) were thawed as per manufacturer’s instructions. Chem-1 cells were loaded with 1 µM Fura-2AM supplemented with 0.01% pluronic F-127 (wt/vol, Life Technologies) in a physiological Ringer’s solution containing (in mM) 140 NaCl, 5 KCl, 10 HEPES, 2 CaCl2, 2 MgCl2 and 10 D-(+)-glucose, pH 7.4 and calcium imaging was performed as described above. Responding cells were defined as those having a > 10% increase from baseline ratio.

#### Whole-cell patch-clamp electrophysiology

HEK293T cells were transiently transfected with Lipofectamine 2000 (11668030, Invitrogen) using (per well): 150 ng human TRPA1 plasmid, 50 ng mCherry plasmid (as a marker for transfection), and 800 ng pcDNA3.1 plasmid. Whole-cell patch-clamp recordings were performed 24-48 hours following transfection. On the day of recordings, cells were detached with TrypLE Express (12604013, Gibco) and seeded onto #1.5 German glass coverslips (502041, World Precision Instruments) at least 3 h before experiments. Whole-cell voltage-clamp recordings were performed with patch pipettes pulled from standard borosilicate glass (1B150F-4, World Precision Instruments) with a P-1000 puller (Sutter Instruments). Patch pipettes had resistances of 2–5 MΩ when filled with internal solution containing (in mM): 140 KCl, 2 MgCl_2_, 1 EGTA, 5 HEPES, and 1 Na_2_ATP, supplemented with 100 µM Na_2_GTP and 100 µM cAMP (pH 7.2, 270 mOsm/kg). Seals and whole-cell configuration were obtained in an external solution containing (in mM): 130 NaCl, 3 KCl, 2.5 CaCl_2_, 1 MgCl_2_, 10 HEPES, and 10 glucose (pH 7.3, 275 mOsm/kg). The pipette offset potential was zeroed before seal formation and whole-cell capacitance was compensated before recording. Current recordings were filtered at 1 kHz (Axopatch 200B, Molecular Devices) and collected at 10 kHz (Digidata 1550B, pCLAMP software, Molecular Devices). The amplifier output gain was set to 2 for all recordings. Whole-cell TRPA1 currents were elicited by applying 300-ms voltage ramps from −100 mV to +100 mV from a holding potential of −20 mV, applied every 3 seconds. All cells had a series resistance of ≤10 MΩ.

#### PAR cleavage assay

Luciferase-based PAR cleavage assays were performed as previously described.^21,64^ Chinese hamster ovary (CHO-K1) cells stably expressing hPAR1, hPAR2, or hPAR4 tagged with nano-luciferase (nLuc) at the N-terminus and enhanced yellow fluorescent protein (eYFP) at the C-terminus were grown. For the cleavage assay, cells were washed with sterile HBSS, and supernatant was collected to measure background luminescence. Cells were incubated for 15 minutes with 100 nM SARS-CoV-2 PLpro, or different concentrations of standard proteases (thrombin 0.001:10 U/ml for PAR1 and 4, and trypsin 0.03:30 nM for PAR2), after which supernatant was collected, mixed with 2 μL/mL Nano-Glo luciferase Assay Substrate Furimazine (Promega, N113A). Luminescence was measured on a Mithras LB 940 Berthold Technologies plate reader with 1 second/well integration time.

#### Neuropeptide release

TG from 7-8 week old mice were cultured overnight as described above and supplemented with 50 ng/mL of nerve growth factor. Cultured neurons were incubated with vehicle or PLpro for 15 minutes in Ringer’s solution at room temperature. For ex vivo trachea stimulation, tracheas were dissected and washed in HBSS for 30 minutes at 37°C. Tracheas were then incubated in DMEM for 30 minutes at 37°C with gentle rotation for baseline measurement and in DMEM with PLpro or vehicle for 1 hour at 37°C with gentle rotation. After incubation, the supernatant (from *in vitro* experiments) and medium (from *ex vivo* experiments) was collected and used to quantify the concentration of neuropeptide using an enzyme linked immunosorbent kit (CGRP: Cayman Chemicals #589001, substance P: Cayman Chemicals #583751) according to the manufacturer’s instructions.

#### Human airway epithelial cell and primary mouse neuron infection of SARS-CoV-2

4-5 x 10^5^ Calu-3 cells were incubated with SARS-CoV-2 (WA1 / USA) in medium (10% heat-inactivated FBS, sodium pyruvate, HEPES, L-glutamine, penicillin/streptomycin) diluted in phenol-red free RPMI to an MOI of 1 for 24 or 48 hours. Mouse TG were cultured as described above and plated in a 96-well plate. TGs (2 each) across 4 animals were pooled and plated across 8 wells. TG cultures were infected with 5 x 10^5^ SARS-CoV-2 virions or medium (mock infection) for 48 hours. Supernatant from SARS-CoV-2- and mock-infected Calu-3 cells was mixed with the PLpro substrate ubiquitin-Rh110 (R&D Systems, #U-555) at a concentration of 670 nM or 1 µM in phenol-free RPMI. Fluorescence (excitation 485nm, emission 535nm) was read on a Spectramax L (Molecular Devices). Samples were run in duplicate. A standard curve was generated by assessing protease activity from a dose-response curve of recombinant SARS-CoV-2 PLpro (R&D), diluted in mock infection medium and measured in parallel. Cell viability was measured using the CellTiter-Glo 2.0 reagent (Promega), prepared according to the manufacturer’s instructions. After removing the supernatant, the reagent was added to the cells, and luminescence was measured on a SpectraMax L. Cell viability was determined by comparing ATP levels to those in mock-infected cells. RNA was isolated with DNA/RNA Shield (Zymo Research) according to the manufacturer’s instructions after removing supernatant from infected cells. RNA library preparation (lnc-RNA library with rRNA removal) and sequencing on an Illumina NovaSeq 6000 were performed by Novogene.

#### Mouse-adapted SARS-CoV-2 infection

Mouse-adapted MA10 strain of SARS-CoV-2 was obtained from Dr. Ralph S. Baric at the University of North Carolina, Chapel Hill. ^65,66^ 8-10 week old C57BL/6 male mice were anesthetized using isoflurane and intranasally inoculated with the MA10 strain at a dose of 10,000 plaque forming units (PFU).

#### MHV-A59 infection

MHV-A59 (ATCC, VR-764) was propagated in NCTC clone 1469 cells (ATCC, CCL-9.1), purified by ultracentrifugation through a 20% sucrose cushion and stored at -80°C. Mice were anesthetized with ketamine and xylazine (80 and 16 mg/kg i.p.) and intranasally inoculated with 40 μL of MHV-A59 at a dose of 25,000 PFU.Viral titers were determined by plaque assay using NCTC clone 929 cells (ATCC, CCL-1). Serial dilutions of viral stocks or bronchoalveolar lavage (BAL) fluid were adsorbed onto cells for 1 h at 37°C. Cells were overlaid with 1.5% agarose-containing plaque assay medium and incubated for 48 h. Plaques were visualized with 0.04% neutral red and counted to determine viral titers (PFU/mL). Bronchoalveolar lavage (BAL) was performed through a tracheal cannula with three 500-μL washes of PBS containing 0.1% BSA and protease inhibitors. BAL fluid was centrifuged at 500 × g for 10 minutes at 4°C, and supernatants were stored at −80°C. Viral titers in BAL fluid were determined by plaque assay.

#### Intranasal behavior

For intranasal behavioral experiments, mice were acclimated for 10 minutes in a custom behavioral chamber (4” x 2” x 6”) with a filter paper bottom (VWR #28321-113) prior to treatment. Mice were treated intranasally with vehicle or PLpro with 10 mg/mL Evan’s blue dye in PBS administered 5µL per nostril (10 µL total) with a P20 pipette. Mice were then placed back in the chamber for recording. Video and audio recordings were performed with a GoPro Hero 11 camera (recording at 2.7 megapixels, 240 Hz) and an aTTo digital audio recorder (recording at 16 kHz). Experimenters were blinded to genotype or treatment. Behavior was scored for 2 minutes by individuals blinded to both genotype and treatment. A nose rub was defined as a singular elliptical stroke made by both forepaws simultaneously, directed to the nose. A face wipe was also a singular elliptical stroke of the forelimbs but contacted the face above the nose and below the ears, making a circular motion. The time of each individual nose rub and face wipe was recorded. A bout of body licking was defined as an uninterrupted series of licking the flank, chest, stomach, and tail. The filter paper that was on the bottom of the chamber was scanned and quantified using *Fiji* by measuring the mean intensity. This mean intensity was inverted (subtracted by 255) and scaled (by 10^8^). No significant difference in sneeze behavior was observed between male and female mice (Figure S11D); therefore, data from both sexes were combined.

#### Whole body plethysmography

Respiratory traces were recorded using Buxco Small Animal whole body plethysmography (WBP) System (DSI, 601-1400-001) and data were collected on the FinePointe Software (DSI). For recordings following acute intranasal treatment, mice were acclimated in the chambers 10 minutes prior to intranasal treatment. Mice were treated intranasally as described above and returned to the WBP chambers for recording. For MHV-A59 and MA10 SARS-CoV-2 infection, mice were recorded for 15 min.

#### Cheek behavior

Itch and acute pain behavioral measurements were performed as described previously.^62^ In brief, mice were shaved and singly housed one week prior to the experiment. Mice were acclimated to the chamber for an hour on both the day prior and day of injection. Compounds were injected intradermally (20 µL) into the cheek. Behavior was videotaped for one hour and scored for either the first 30 minutes (scratching) or the first 5 minutes (wiping) as previously described.^67^

#### Von Frey and radiant heat behavior

Mice were lightly anesthetized with isoflurane (2.5%) and received intradermal injections (15 µL) of PLpro or vehicle into the plantar surface of the hind paw. For von Frey, both hind paws were injected, one with treatment and the other with vehicle solution. Mice were acclimated in behavioral chambers for two consecutive days for at least one hour each day, followed by an additional 30 minute acclimation with the investigator in the room. Mechanical threshold was measured using calibrated von Frey monofilaments (Touch Test) on a metal grate platform (IITC). Von Frey was performed as previously described^68,69^ using the up-down method.^70^ Measurements were taken one day prior to injection, and 3 and 24 hours after injection. Radiant heat assay was performed using the Hargreaves test system (IITC Life Science) as previously described.^68,71^ A glass platform was set at 30°C during the acclimation and throughout the experiment and the radiant heat source raised the platform temperature to 41.5°C within 5 s and to 44.9°C within 10 s, as measured by a fast temperature probe (Physitemp). Valid responses for both von Frey and radiant heat included fast paw withdrawal, licking/biting/shaking of the affected paw, or flinching. Experimenter was blinded to treatment and genotype.

### Quantification and Statistical Analysis

#### *In vivo* imaging analysis

Motion correction and source extraction were performed using Suite2p.^72^ Suite2p-generated regions of interest (ROIs) were manually inspected and additional ROIs were manually identified. Neuropil was subtracted with a coefficient of 0.7. F_0_ is defined as the median fluorescence 150 frames prior to stimulus treatment and used to calculate dF/F_0_. Normalized traces were Gaussian filtered (σ = 2 frames) for stimulus-response analysis and separately filtered (σ = 3 frames) for identification of spontaneous baseline activity. Cells were identified as spontaneously active if peak fluorescence during the baseline window > 5% of the median fluorescence. Stimulus responders were identified if the peak normalized fluorescence during the 2-minute stimulation period increased by at least 10% relative to the median normalized fluorescence during the preceding 150 frames. Cells were identified as spontaneously active if peak fluorescence during the baseline window > 5% of the median fluorescence. The percentage of responsive cells was calculated by dividing the number of neurons responding to a given stimulus by the total number of neurons that responded to at least one intranasally applied stimulus. Neurons with spontaneous baseline activity were excluded.

#### *In vitro* imaging analysis

Responding neurons were defined as those having a > 10% increase from baseline ratio. Human iPSC-derived neurons were classified as responsive when the ratio increased by > 5% from baseline. All calcium imaging analyses were performed using custom Python scripts. Each biological replicate (mice unless noted otherwise) represents an average of 3-4 wells with ∼100-300 neurons per well. Control and experimental groups within each figure were collected concurrently and thus, each comparison is internally matched and not affected by between-experiment variation.

#### Audio and video sneeze analysis

Sneeze events were identified from peaks in the audio recordings. Audio recordings were cropped to the start of the experiment and a 2-3 second recording of background sound from the recording was used for noise reduction in Audacity (Noise reduction: 30 dB, sensitivity: 6, frequency of smoothing: 6 bands). Denoised audio recordings were then analyzed using a custom Python script. Audio peaks were identified using the SciPy “find_peaks” function (peak prominence > 1000 and a minimum distance of 0.2 s between peaks). The audio recordings from inside the chamber were aligned to the video and audio recordings from the GoPro (outside the chamber). Audio peaks recorded both inside and outside the chamber were excluded as background noise. Audio peaks that correlated with other behaviors such as biting the chamber and body licking were excluded.

#### Whole body plethysmography sneeze and cough analysis

Box flow was analyzed using custom Python scripts and aligned with video recordings to identify instances of dye expulsion. Peaks ≥ 1.10-fold greater than the mean of the three preceding peaks were identified and manually classified as coughs or one of two sneeze waveforms. Coughs were characterized by an extended inspiratory peak preceding an expiratory peak, whereas sneezes were characterized by either a larger expiratory peak or two sharp expiratory peaks in succession.

#### RNA sequencing analysis

For quantification of viral transcripts, reads were mapped to the appropriate host (human or mouse) transcriptome concatenated with the SARS-CoV-2 transcriptome using *Kallisto (0.44.0)*. For analysis of infected mouse TG, reads were mapped to the mouse genome concatenated with the SARS-CoV-2 genome using *STAR (2.7.10a)*. Gene counts were extracted using featureCounts in *Subread (2.0.1).* All but one sample contained more than 80% mapped features; one sample contained 68% mapped reads and was excluded based on the robust principal component analysis method.^73^ The uniquely aligned features were used for differential expression analysis using *DESeq2.* Gene sets associated with functional neuronal subtypes were from a previous study.^57^ As performed in the paper, we pooled the identical Gene Ontology terms Mouse Genome Informatics (MGI) (updated as of November 2023) except for the itch gene list, which we used from the paper. To determine whether these custom gene sets exhibited greater differential expression than expected by chance, we performed an empirical resampling test. For each gene set, we generated a null distribution by randomly sampling 100,000 gene sets of equal size from all transcripts detected across the samples and calculating the median absolute log₂ fold change for each sampled set. The empirical *p* value was calculated as the proportion of sampled gene sets with a median absolute log₂ fold change equal to or greater than that of the observed gene set.

#### Statistical Analysis

All statistical tests were two-tailed where applicable. Sample sizes (n), test statistics, exact p values, and degrees of freedom (df) are reported, where relevant, in the figure legends. No statistical methods were used to predetermine sample sizes. Experiments comparing genotypes were performed and analyzed with investigators blinded to animal genotype. All experiments were independently repeated at least twice, and data from independent experiments were pooled. Graphing and analyses were performed in GraphPad Prism 11 (GraphPad Software) or using custom scripts in Python.

## Key resources table

| REAGENT or RESOURCE | SOURCE | IDENTIFIER |
|---|---|---|
| Bacterial and virus strains |  |  |
| SARS-CoV-2 (USA-WA1/2020) | BEI Resource | NR-52281 |
| Mouse-adapted SARS-CoV-2 (MA10) | Leist et al.<sup>66</sup> | N/A |
| Murine Hepatitis Virus (MHV-A59) | ATCC | ATCC VR-764 |
| Chemicals, peptides, and recombinant proteins |  |  |
| SARS-CoV-2 PLpro | R&D Systems | Cat#E-611 |
| SARS-CoV-2 PLpro | Cayman Chemical | Cat#31817 |
| SARS-CoV-2 PLpro C111S mutant | R&D Systems | Custom order |
| SARS-CoV PLpro | R&D Systems | Cat#E-610 |
| MERS-CoV PLpro | R&D Systems | Cat#E-609 |
| SARS-CoV-2 Mpro (3CLpro) | R&D Systems | Cat#E-720 |
| (*E*)-Capsaicin | Tocris | Cat#0462 |
| Allyl isothiocyanate (AITC) | Sigma Aldrich | Cat#377430 |
| HC-030031 | Tocris | Cat#2896 |
| AMG 517 | Sigma Aldrich | Cat#SML2717 |
| PLpro inhibitor 19 | Tocris | Cat#7505 |
| Fura-2 AM | Thermo fisher | Cat#F1201 |
| Pluronic F-127 | Thermo Scientific | Cat#P6867 |
| Evan’s blue dye | Sigma Aldrich | Cat#E2129 |
| Nerve growth factor | Invitrogen | Cat#A42627 |
| Critical commercial assays |  |  |
| CGRP ELISA kit | Cayman Chemical | Cat#589001 |
| Substance P ELISA kit | Cayman Chemical | Cat#583751 |
| CellTiter-Glo 2.0 | Promega | Cat#G9241 |
| DNA/RNA Shield | Zymo Research | Cat#R1100 |
| Nano-Glo Luciferase Assay Substrate (furimazine) | Promega | Cat#N113A |
| Ubiquitin-Rh110 | R&D Systems | Cat#U-555 |
| Deposited data |  |  |
| Bulk RNA-seq of SARS-CoV-2-infected and mock-infected human Calu-3 cells and mouse trigeminal ganglion cultures | This study | GEO: GSE252056<br>https://www.ncbi.nlm.nih.gov/geo/query/acc.cgi?acc=GSE252056 |
| Experimental models: Cell lines |  |  |
| HEK293T | ATCC | CRL-3216: RRID: CVCL_0063 |
| CHO-K1 | ATCC | CCL-61; RRID: CVCL_0214 |
| Calu-3 | ATCC | HTB-55 |
| NCTC clone 1469 | ATCC | CCL-9.1; RRID: CVCL_3066 |
| NCTC clone 929 | ATCC | CCL-1; RRID: CVCL_0462 |
| Chem-1 (MRGPRX1-expressing) | Eurofins DiscoverX | HTS122RTA |
| Chem-1 (MRGPRX2-expressing) | Eurofins DiscoverX | HTS058RTA |
| Human iPSC-derived neurons | Axol Bioscience | ax0555 |
| Experimental models: Organisms/strains |  |  |
| C57BL/6J | Jackson Laboratories | JAX:000664; RRID: IMSR_JAX:000664 |
| C57BL/6NCrl | Charles River | CR:027; RRID: IMSR_CRL:027 |
| *Pirt*<sup>Cre</sup>: Pirttm3.1(cre)Xzd | Kim et al.<sup>74</sup> | MGI:6332952 |
| Ai96 | Jackson Laboratories | JAX: 028866; RRID: IMSR_JAX:028866 |
| *Trpv1* KO: Trpv1tm1Jul | Jackson Laboratories | JAX: 003770; RRID: IMSR_JAX:003770 |
| *Trpa1* KO: Trpa1tm1Jul | Bautista et al.<sup>40</sup> | MGI:3692770 |
| *Trpa1* flox: B6.129S-Trpa1tm2Kykw/J | Jackson Laboratories | JAX: 008650; RRID: IMSR_JAX:008650 |
| *Scn10a*<sup>Cre</sup>: B6.129(Cg)-Scn10atm2(cre)Jwo/TjpJ | Jackson Laboratories | JAX: 036564; RRID: IMSR_JAX:036564 |
| Recombinant DNA |  |  |
| human TRPA1 | Wilson et al.<sup>62</sup> | N/A |
| Software and algorithms |  |  |
| Prism 11 | GraphPad Software | RRID: SCR_002798 |
| Fiji (ImageJ) | Schindelin et al.<sup>75</sup> | http://fiji.sc/<br>RRID: SCR_002285 |
| pClamp | Molecular Devices | RRID: SCR_011323 |
| FinePointe Software | Data Sciences International | RRID: SCR_028683 |
| Python version 3.10 | Python Software Foundation | RRID: SCR_008394 |
| Suite2P | Pachitariu et al.<sup>72</sup> | https://github.com/cortex-lab/Suite2P<br>RRID: SCR_016434 |
| SciPy | Virtanen et al.<sup>76</sup> | http://www.scipy.org/<br>RRID: SCR_008058 |
| Audacity | Audacity Team | http://audacity.sourceforge.net/<br>RRID: SCR_007198 |
| Adobe Premiere Pro | Adobe Creative Cloud | RRID: SCR_021315 |
| Kallisto (0.44.0) | Bray et al.<sup>77</sup> | https://pachterlab.github.io/kallisto/about RRID: SCR_016582 |
| STAR (2.7.10a) | Dobin et al.<sup>78</sup> | https://code.google.com/p/rna-star/<br>RRID: SCR_004463 |
| Subread (2.0.1) | Liao et al.<sup>79</sup> | http://subread.sourceforge.net/<br>RRID: SCR_009803 |
| DESeq2 | Love et al.<sup>80</sup> | https://bioconductor.org/packages/release/bioc/html/DESeq2.html RRID: SCR_015687 |
| Custom code | This paper | Zenodo: 10.5281/zenodo.23147030 |
| Other |  |  |
| Buxco Small Animal Whole Body Plethysmography (WBP) System | Data Sciences International | Cat#601-1400-001 |
| Von Frey filaments | North Coast | Cat#NC12775 |
| Hargreaves test system | IITC | Cat#390G |
| Filter paper | VWR | Cat#28321-113 |
| GoPro Hero 11 | GoPro | N/A |
| aTTo digital audio recorder | aTTo | N/A |

## ACKNOWLEDGEMENTS

We are grateful for Dr. Sven-Eric Jordt, Dr. Anabel De Caceres Bustos, Odilia Liu, and Abigail Mende for critical discussions and assistance with pilot experiments. We thank Dr. Rachel Brem, Dr. Jay Parrish, and the 2022 MBL Neurobiology Course students for assistance with pilot RNA-seq analysis; Dr. Olivia Goldman, Dr. Erin Aisenberg and Lillian Murphy for critical review of the manuscript; and Dr. Julia Bleier for PDB structure visualization. During the preparation of this work, the authors used ChatGPT (OpenAI) and Claude (Anthropic) to assist with formatting and improving clarity. After using these tools, the authors carefully reviewed and edited the content and take full responsibility for the content of the publication.

## AUTHOR CONTRIBUTIONS

Conceptualization: S.S.M. and D.M.B. Methodology: S.S.M., Z.G., R.S., A.N.F., Y.M. Investigation: S.S.M., Z.G., R.S., A.N.F., C.P.C., L.S., J.S., M.C., C.J.A., A.A.K.M., C.V. Formal analysis: S.S.M., Z.G., R.S., A.N.F., J.S., C.J.A., A.A.K.M., U.V., D.M.B. Visualization: S.S.M and D.M.B. Supervision: J.K., J.S.C., R.R., S.A.S., D.M.B. Project administration: S.S.M. and D.M.B. Funding acquisition: D.M.B. writing – original draft: S.S.M. and D.M.B. writing – review & editing: S.S.M. and D.M.B. with input from all authors.

## DATA AND CODE AVAILABILITY

RNA-seq data generated in this study have been deposited in the Gene Expression Omnibus (GEO) under accession number GSE252056. All original code has been deposited to Zenodo and is publicly available at <u>10.5281/zenodo.23147030</u>. This study did not generate new unique reagents. Further information and requests for resources and reagents should be directed to and will be fulfilled by the Lead Contact, Diana M. Bautista.

## REFERENCES

1. Bliddal, S. et al. Acute and persistent symptoms in non-hospitalized PCR-confirmed COVID-19 patients. Sci. Rep. 11, 13153 (2021).

2. Grant, M. C. et al. The prevalence of symptoms in 24,410 adults infected by the novel coronavirus (SARS-CoV-2; COVID-19): A systematic review and meta-analysis of 148 studies from 9 countries. PloS One 15, (2020).

3. Jiang, H. et al. Divergent sensory pathways of sneezing and coughing. Cell 187, 5981–5997 14 (2024).

4. Pinho-Ribeiro, F. A., Verri, W. A. & Chiu, I. M. Nociceptor Sensory Neuron–Immune Interactions in Pain and Inflammation. Trends Immunol. 38, 5–19 (2017).

5. Prescott, S. L., Umans, B. D., Williams, E. K., Brust, R. D. & Liberles, S. D. An Airway Protection Program Revealed by Sweeping Genetic Control of Vagal Afferents. Cell 181, 574–589.e14 (2020).

6. Gannot, N. et al. A vagal–brainstem interoceptive circuit for cough-like defensive behaviors in mice. Nat. Neurosci. 1–11, (2024).

7. Prescott, S. L. & Liberles, S. D. Internal senses of the vagus nerve. Neuron 110, 579–599 (2022).

8. Staurengo-Ferrari, L., Deng, L. & Chiu, I. M. Interactions between nociceptor sensory neurons and microbial pathogens in pain. PAIN 163, S57 (2022).

9. Akira, S., Uematsu, S. & Takeuchi, O. Pathogen recognition and innate immunity. Cell 124, 783–801 (2006).

10. Hartenian, E. et al. The molecular virology of Coronaviruses. J. Biol. Chem. 10.1074/jbc.REV120.013930 (2020) doi:10.1074/jbc.REV120.013930.

11. Shin, D. et al. Papain-like protease regulates SARS-CoV-2 viral spread and innate immunity. Nature 587, 657–662 (2020).

12. Reddy, V. B., Iuga, A. O., Shimada, S. G., LaMotte, R. H. & Lerner, E. A. Cowhage-evoked itch is mediated by a novel cysteine protease: a ligand of protease-activated receptors. J. Neurosci. Off. J. Soc. Neurosci. 28, 4331–4335 (2008).

13. Kwong, K., Nassenstein, C., Garavilla, L., Meeker, S. & Undem, B. J. Thrombin and trypsin directly activate vagal C-fibres in mouse lung via protease-activated receptor-1. J. Physiol. 588, 1171–1177 (2010).

14. Liu, Q. et al. The Distinct Roles of Two GPCRs, MrgprC11 and PAR2, in Itch and Hyperalgesia. Sci. Signal. 4, ra45–ra45 (2011).

15. Gu, Q. & Lee, L.-Y. House dust mite potentiates capsaicin-evoked Ca2+ transients in mouse pulmonary sensory neurons via activation of protease-activated receptor-2. Exp. Physiol. 97, 534–543 (2012).

16. Jimenez-Vargas, N. N. et al. Protease-activated receptor-2 in endosomes signals persistent pain of irritable bowel syndrome. Proc. Natl. Acad. Sci. U. S. A. 115, 7438–7447 (2018).

17. Serhan, N. et al. House dust mites activate nociceptor-mast cell clusters to drive type 2 skin inflammation. Nat. Immunol. 20, 1435–1443 (2019).

18. Perner, C. et al. Substance P Release by Sensory Neurons Triggers Dendritic Cell Migration and Initiates the Type-2 Immune Response to Allergens. Immunity 53, 1063–1077 7 (2020).

19. Hassler, S. N., et al. The cellular basis of protease-activated receptor 2-evoked mechanical and affective pain. JCI Insight 5, (2020).

20. Pradhananga, S., Tashtush, A. A., Allen-Vercoe, E., Petrof, E. O. & Lomax, A. E. Protease-dependent excitation of nodose ganglion neurons by commensal gut bacteria. J. Physiol. 598, 2137–2151 (2020).

21. Deng, L. et al. Staphylococcus aureus Drives Itch and Scratch-Induced Skin Damage Through a V8 Protease-PAR1 Axis. Cell (2023) 10.1016/j.cell.2023.10.019.

22. Liang, Q. et al. An electrochemical biosensor for SARS-CoV-2 detection via its papain-like cysteine protease and the protease inhibitor screening. Chem. Eng. J. 10.1016/j.cej.2022.139646 (2023) doi:10.1016/j.cej.2022.139646.

23. Jimenez-Campos, A. G. et al. A cell-based Papain-like Protease (PLpro) activity assay for rapid detection of active SARS-CoV-2 infections and antivirals. PloS One 19, (2024).

24. Shusterman, D. Trigeminal Function in Sino-Nasal Health and Disease. Biomedicines 11, 1778 (2023).

25. Sekizawa, S. & Tsubone, H. Nasal mechanoreceptors in guinea pigs. Respir. Physiol. 106, 223–230 (1996).

26. Tsubone, H. Nasal ‘pressure’ receptors. *Nihon Juigaku Zasshi Jpn*. J. Vet. Sci. 52, 225–232 (1990).

27. Stirling, L. C. et al. Nociceptor-specific gene deletion using heterozygous NaV1.8-Cre recombinase mice. Pain 113, 27–36 (2005).

28. Baraniuk, J. N. & Kim, D. Nasonasal reflexes, the nasal cycle, and sneeze. Curr. Allergy Asthma Rep. 7, 105–111 (2007).

29. Li, F. et al. Sneezing reflex is mediated by a peptidergic pathway from nose to brainstem. Cell 184, 3762–3773 10 (2021).

30. Strassman, A. M. & Levy, D. Response properties of dural nociceptors in relation to headache. J. Neurophysiol. 95, 1298–1306 (2006).

31. Canning, B. J. et al. Identification of the tracheal and laryngeal afferent neurones mediating cough in anaesthetized guinea-pigs. J. Physiol. 557, 543–558 (2004).

32. Zhang, C., Lin, R.-L., Hong, J., Khosravi, M. & Lee, L.-Y. Cough and expiration reflexes elicited by inhaled irritant gases are intensified in ovalbumin-sensitized mice. Am. J. Physiol. Regul. Integr. Comp. Physiol. 312, 718–726 (2017).

33. Sun, H. et al. KCNQ/M-channels regulate mouse vagal bronchopulmonary C-fiber excitability and cough sensitivity. JCI Insight 4, (2019).

34. Iwasaki, N. et al. Allergen endotoxins induce T-cell-dependent and non-IgE-mediated nasal hypersensitivity in mice. J. Allergy Clin. Immunol. 139, 258–268 10 (2017).

35. Kayasuga, R., Sugimoto, Y., Watanabe, T. & Kamei, C. Histamine H1 receptors are involved in mouse nasal allergic responses: a demonstration with H1 receptor-deficient mice. Int. Immunopharmacol. 2, 745–750 (2002).

36. Báez-Santos, Y. M., Mielech, A. M., Deng, X., Baker, S. & Mesecar, A. D. Catalytic function and substrate specificity of the papain-like protease domain of nsp3 from the Middle East respiratory syndrome coronavirus. J. Virol. 88, 12511–12527 (2014).

37. Osipiuk, J. et al. Structure of papain-like protease from SARS-CoV-2 and its complexes with non-covalent inhibitors. Nat. Commun. 12, 743 (2021).

38. Han, Y.-S. et al. Papain-like protease 2 (PLP2) from severe acute respiratory syndrome coronavirus (SARS-CoV): expression, purification, characterization, and inhibition. Biochemistry 44, 10349–10359 (2005).

39. Kanjanahaluethai, A., Jukneliene, D. & Baker, S. C. Identification of the murine coronavirus MP1 cleavage site recognized by papain-like proteinase 2. J. Virol. 77, 7376–7382 (2003).

40. Bautista, D. M. et al. TRPA1 mediates the inflammatory actions of environmental irritants and proalgesic agents. Cell 124, 1269–1282 (2006).

41. Grace, M., Birrell, M. A., Dubuis, E., Maher, S. A. & Belvisi, M. G. Transient receptor potential channels mediate the tussive response to prostaglandin E2 and bradykinin. Thorax 67, 891–900 (2012).

42. Hill, R. Z., Morita, T., Brem, R. B. & Bautista, D. M. S1PR3 Mediates Itch and Pain via Distinct TRP Channel-Dependent Pathways. J. Neurosci. Off. J. Soc. Neurosci. 38, 7833–7843 (2018).

43. Chiu, I. M. et al. Bacteria activate sensory neurons that modulate pain and inflammation. Nature 501, 52–57 (2013).

44. Wang, H. & Woolf, C. J. Pain TRPs. Neuron 46, 9–12 (2005).

45. Grace, M. S. & Belvisi, M. G. TRPA1 receptors in cough. Pulm. Pharmacol. Ther. 24, 286–288 (2011).

46. Serafini, R. A. et al. SARS-CoV-2 airway infection results in the development of somatosensory abnormalities in a hamster model. Sci. Signal. 16, eade4984 (2023).

47. Sudre, C. H. et al. Attributes and predictors of long COVID. Nat. Med. 27, 626–631 (2021).

48. Caterina, M. J. et al. Impaired nociception and pain sensation in mice lacking the capsaicin receptor. Science 288, 306–313 (2000).

49. Lennertz, R. C., Kossyreva, E. A., Smith, A. K. & Stucky, C. L. TRPA1 mediates mechanical sensitization in nociceptors during inflammation. PloS One 7, (2012).

50. Basbaum, A. I., Bautista, D. M., Scherrer, G. & Julius, D. Cellular and molecular mechanisms of pain. Cell 139, 267–284 (2009).

51. Tamari, M. et al. Sensory neurons promote immune homeostasis in the lung. Cell 187, 44–61.e17 (2024).

52. Barbosa Bomfim, C. C., et al. CGRP inhibits SARS-CoV-2 infection of bronchial epithelial cells, and its pulmonary levels correlate with viral clearance in critical COVID-19 patients. J. Virol. 98, (2024).

53. Joyce, J. D., et al. SARS-CoV-2 rapidly infects peripheral sensory and autonomic neurons, contributing to central nervous system neuroinvasion before viremia. Int. J. Mol. Sci. 25, 8245 (2024).

54. Meinhardt, J. et al. Olfactory transmucosal SARS-CoV-2 invasion as a port of central nervous system entry in individuals with COVID-19. Nat. Neurosci. 10.1038/s41593-020-00758-5 (2020) doi:10.1038/s41593-020-00758-5.

55. Flamier, A., Bisht, P., Richards, A., Tomasello, D. L. & Jaenisch, R. Human iPS cell-derived sensory neurons can be infected by SARS-CoV-2. (2023) doi:10.1016/j.isci.2023.107690.

56. Co, V. A. et al. SARS-CoV-2 infects olfactory neurons and basal stem cells and induces axonal degeneration through TRPV1 activation. iScience 29, 116098 (2026).

57. Usoskin, D. et al. Unbiased classification of sensory neuron types by large-scale single-cell RNA sequencing. Nat. Neurosci. 18, 145–153 (2015).

58. Ruhl, C. R. et al. Mycobacterium tuberculosis Sulfolipid-1 Activates Nociceptive Neurons and Induces Cough. Cell 181, 293–305.e11 (2020).

59. Baral, P. et al. Nociceptor sensory neurons suppress neutrophil and γδ T cell responses in bacterial lung infections and lethal pneumonia. Nat. Med. 24, 417–426 (2018).

60. Granton, E. et al. Biofilm exopolysaccharides alter sensory-neuron-mediated sickness during lung infection. Cell 187, 1874–1888.e14 (2024).

61. Kim, J. S., Ru, F., Meeker, S. & Undem, B. J. Direct activation of airway sensory C-fibers by SARS-CoV-2 S1 spike protein. Physiol. Rep. 11, e15900 (2023).

62. Wilson, S. R. et al. TRPA1 is required for histamine-independent, Mas-related G protein–coupled receptor–mediated itch. Nat. Neurosci. 14, 595–602 (2011).

63. Moayedi, Y. The cellular basis of mechanosensation in mammalian tongue. Cell Rep 42, 112087 (2023).

64. Chandrabalan, A. et al. Human osteoarthritis knee joint synovial fluids cleave and activate Proteinase-Activated Receptor (PAR) mediated signaling. Sci. Rep. 13, (2023).

65. Biering, S. B. et al. Screening a Library of FDA-Approved and Bioactive Compounds for Antiviral Activity against SARS-CoV-2. ACS Infect. Dis. 7, 2337–2351 (2021).

66. Leist, S. R. et al. A Mouse-Adapted SARS-CoV-2 Induces Acute Lung Injury and Mortality in Standard Laboratory Mice. Cell 183, 1070–1085.e12 (2020).

67. Shimada, S. G. & LaMotte, R. H. Behavioral differentiation between itch and pain in mouse. Pain 139, 681–687 (2008).

68. Tsunozaki, M. et al. A “toothache tree” alkylamide inhibits Aδ mechanonociceptors to alleviate mechanical pain. J. Physiol. 591, 3325–3340 (2013).

69. Chaplan, S. R., Bach, F. W., Pogrel, J. W., Chung, J. M. & Yaksh, T. L. Quantitative assessment of tactile allodynia in the rat paw. J. Neurosci. Methods 53, 55–63 (1994).

70. Dixon, W. J. The Up-and-Down Method for Small Samples. J. Am. Stat. Assoc. 60, 967–978 (1965).

71. Hargreaves, K., Dubner, R., Brown, F., Flores, C. & Joris, J. A new and sensitive method for measuring thermal nociception in cutaneous hyperalgesia. Pain 32, 77–88 (1988).

72. Pachitariu, M., et al. Suite2p: beyond 10,000 neurons with standard two-photon microscopy. in *In bioRxiv* 061507 (2017). doi:10.1101/061507.

73. Chen, X., Zhang, B., Wang, T., Bonni, A. & Zhao, G. Robust principal component analysis for accurate outlier sample detection in RNA-Seq data. BMC Bioinformatics 21, 269 (2020).

74. Kim, A. Y. et al. Pirt, a phosphoinositide-binding protein, functions as a regulatory subunit of TRPV1. Cell 133, 475–485 (2008).

75. Schindelin, J. et al. Fiji: an open-source platform for biological-image analysis. Nat. Methods 9, 676–682 (2012).

76. Virtanen, P. et al. SciPy 1.0: fundamental algorithms for scientific computing in Python. Nat. Methods 17, 261–272 (2020).

77. Bray, N. L., Pimentel, H., Melsted, P. & Pachter, L. Near-optimal probabilistic RNA-seq quantification. Nat. Biotechnol. 34, 525–527 (2016).

78. Dobin, A. et al. STAR: ultrafast universal RNA-seq aligner. Bioinformatics 29, 15–21 (2013).

79. Liao, Y., Smyth, G. K. & Shi, W. The Subread aligner: fast, accurate and scalable read mapping by seed-and-vote. Nucleic Acids Res. 41, e108–e108 (2013).

80. Love, M. I., Huber, W. & Anders, S. Moderated estimation of fold change and dispersion for RNA-seq data with DESeq2. Genome Biol. 15, 550 (2014).

