## Supplemental Figures and Table for "A coronavirus protease activates sensory neurons via TRPA1 to drive sneeze, cough, and pain"

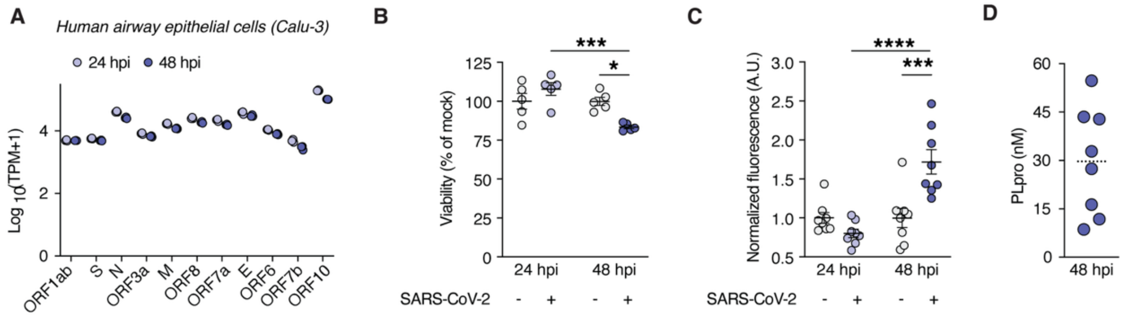


**Figure S1. Extracellular PLpro activity is detected following SARS-CoV-2 infection of airway epithelial cells, related to Figure 1.**

(A) SARS-CoV-2 transcript expression (transcripts per million, TPM) in Calu-3 cells infected at a multiplicity of infection (MOI) of 1, measured at 24 and 48 hours post-infection (hpi). $n$ = 3 wells per condition.

(B) Infection induces cell death at 48 hpi measured as the percentage of viable cells relative to mock-infected controls. Two-way ANOVA**:** $F(1,16) = 11.69$, $p$ (Interaction of time and infection) = 0.0035; Šídák's multiple comparisons**:** $p$ (mock vs. infected 24 hpi) = 0.3628, $p$ (mock vs. infected 48 hpi) = 0.0140, $p$ (24 hpi vs. 48 hpi infected) = 0.0005, $n$ = 5 wells per group. Error bars represent mean ± SEM**.**

(C) Proteolytic activity of PLpro in culture medium from SARS-CoV-2- or mock-infected Calu-3 cells at 24 and 48 hpi, measured by fluorescence of cleaved substrate and normalized to same-day controls. Two-way ANOVA**:** $F(1,28) = 18.01$, $p$ (Interaction of time and infection) = 0.0002; Šídák's multiple comparisons**:** $p$ (mock vs. infected 24 hpi) = 0.5022, $p$ (mock vs. infected 48 hpi) = 0.0002, $p$ (24 hpi vs. 48 hpi infected) < 0.0001, $n$ = 8 wells per group. Data from two independent experiments. Error bars represent mean ± SEM**.**

(D) Interpolated concentration of PLpro in the supernatant of Calu-3 cells at 48 hpi, $n$ = 8 wells; line represents the mean.

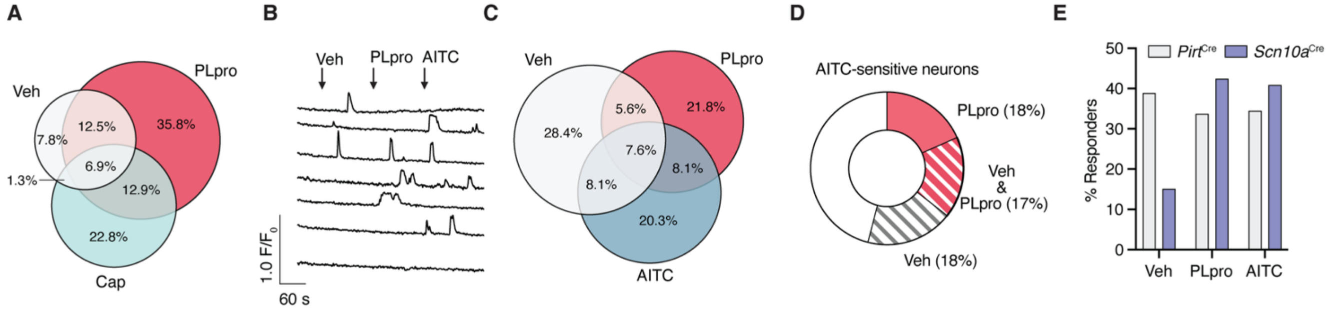


**Figure S2. PLpro activates subsets of nociceptors in vivo, related to Figure 1.**
(A) Venn diagram showing overlap among neurons responsive to vehicle (Veh), PLpro, and capsaicin (Cap). Percentages indicate the proportion of the 232 neurons that responded to at least one stimulus. In total, 66/232 neurons (28.4%) responded to vehicle, 158/232 (68.1%) responded to PLpro, and 102/232 (44.0%) responded to capsaicin.
(B) Representative calcium traces showing responses to sequential administration of vehicle, PLpro, and AITC.
(C) Venn diagram showing overlap among neurons responsive to vehicle, PLpro, and AITC. Percentages indicate the proportion of the 197 neurons that responded to at least one stimulus. In total, 98/197 neurons (49.7%) responded to vehicle, 85/197 (43.1%) responded to PLpro, and 87/197 (44.2%) responded to AITC.
(D) PLpro activates a subset (35.6%, 31/87) of AITC-sensitive neurons; data from 4 mice.
(E) Percentage of recorded neurons responding to vehicle, PLpro, and AITC when GCaMP6s was expressed broadly across peripheral sensory neurons (*Pirt*$^{Cre}$) or in Nav1.8-expressing nociceptors (*Scn10a*$^{Cre}$). *Pirt*$^{Cre}$ data include 252 neurons from 4 mice; *Scn10a*$^{Cre}$ data include 66 neurons from 3 mice.

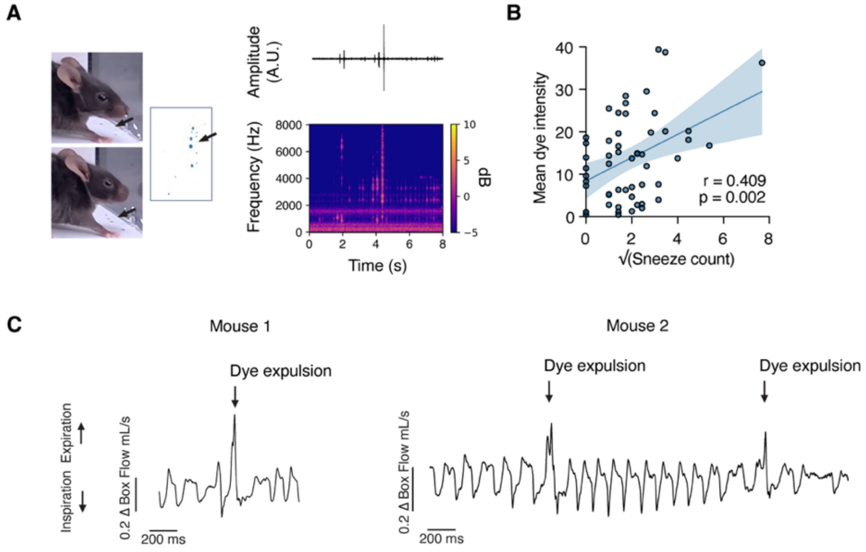


**Figure S3. Nasal dye expulsions correlate with high-frequency audible events and respiratory waveforms, related to Figure 1.**

(A) Sneeze events identified from audio recordings co-occurred with dye expulsion as indicated by dye droplets deposited on the chamber floor. Left, images before and after a dye expulsion event; arrow indicates dye deposited by this sneeze on the bottom of the chamber. Right, audio waveform and spectrogram recorded during the same event.

(B) The number of sneezes over the 2-minute recording period correlates with the intensity of expelled dye on the bottom of the chamber. Square root-transformed sneeze count is significantly correlated with the mean dye intensity (Pearson's r = 0.409, $p$ = 0.0023).

(C) Dye expulsion events co-occur with distinct respiratory waveform patterns. Representative airflow traces (Δ box flow) show a single dye expulsion event in an animal (left), with a waveform resembling a sneeze, and two consecutive dye expulsion events in a different animal (right), with waveforms resembling a sneeze (first event) and a cough (second event), respectively.

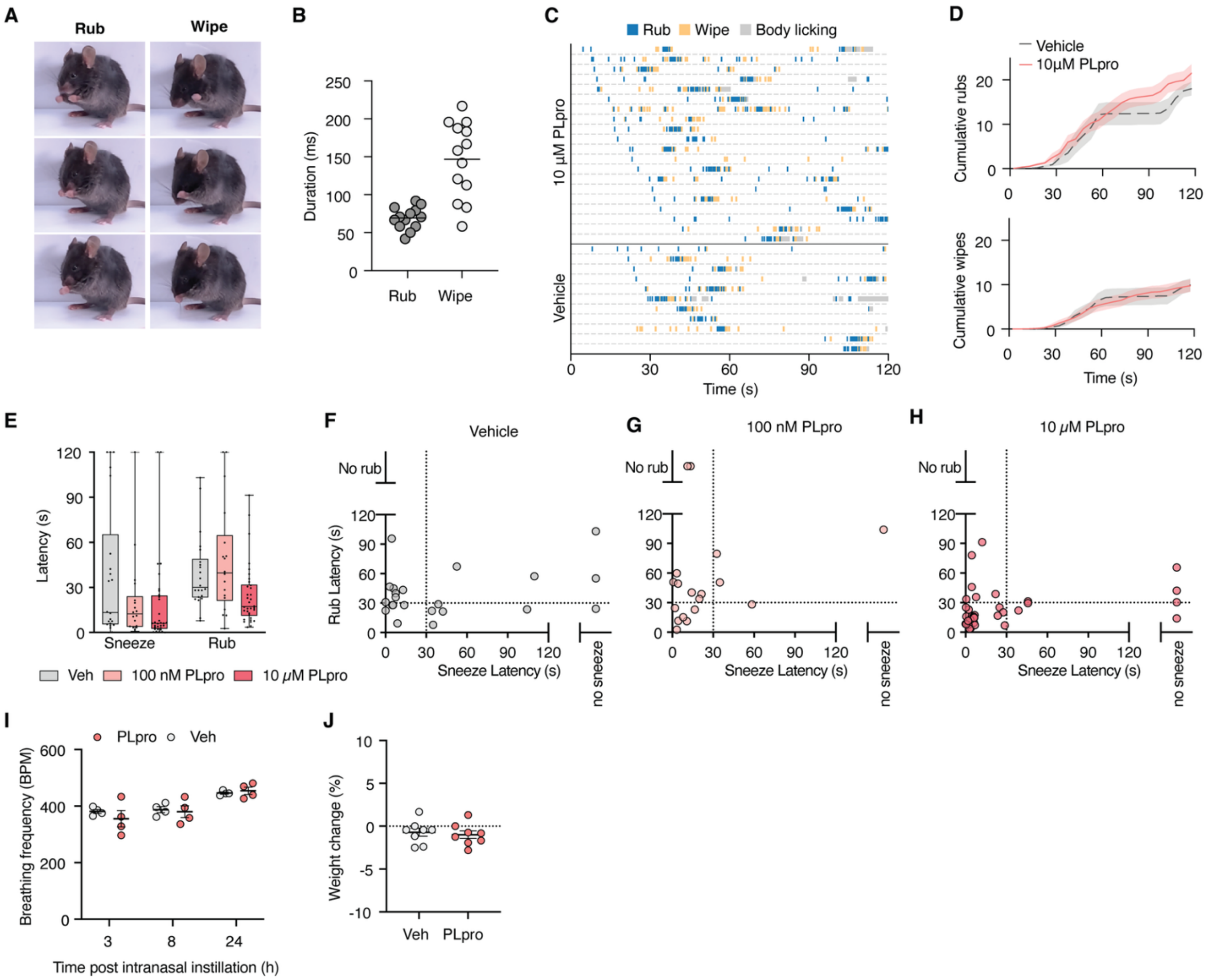


**Figure S4. Characterization of behaviors elicited by intranasal PLpro treatment, related to Figure 1.**

(A) Image sequence of a nose rub and face wipe.

(B) Duration (milliseconds) of the behavioral sequence for a nose rub and face wipe.

(C) Raster plot of rubs, wipes, and body-licking events in mice treated intranasally with 10 μM PLpro or vehicle-treated mice.

(D) Average cumulative rub and wipe counts from individual mice treated with 10 μM PLpro ($n$ = 20) or vehicle ($n$ = 11).

(E) Latencies to the first sneeze and first rub after vehicle, 100 nM PLpro, or 10 μM PLpro.

(F–H) Relationship between sneeze and rub latencies after (F) vehicle ($n$ = 22 mice), (G) 100 nM PLpro ($n$ = 18 mice), or (H) 10 μM PLpro ($n$ = 34 mice).

(I) Intranasal instillation of 40 μL of 12.5 μM PLpro (1.14 μg/μL) does not alter breathing frequency at 3, 8, or 24 h post-instillation, $n$ = 4 mice per group. (**J**) Intranasal instillation of 40 μL of 12.5 μM PLpro (1.14 μg/μL) does not induce weight loss 24 hours post-instillation relative to pretreatment, $n$ = 8 mice per group.

Error bars and shading represent mean ± SEM.

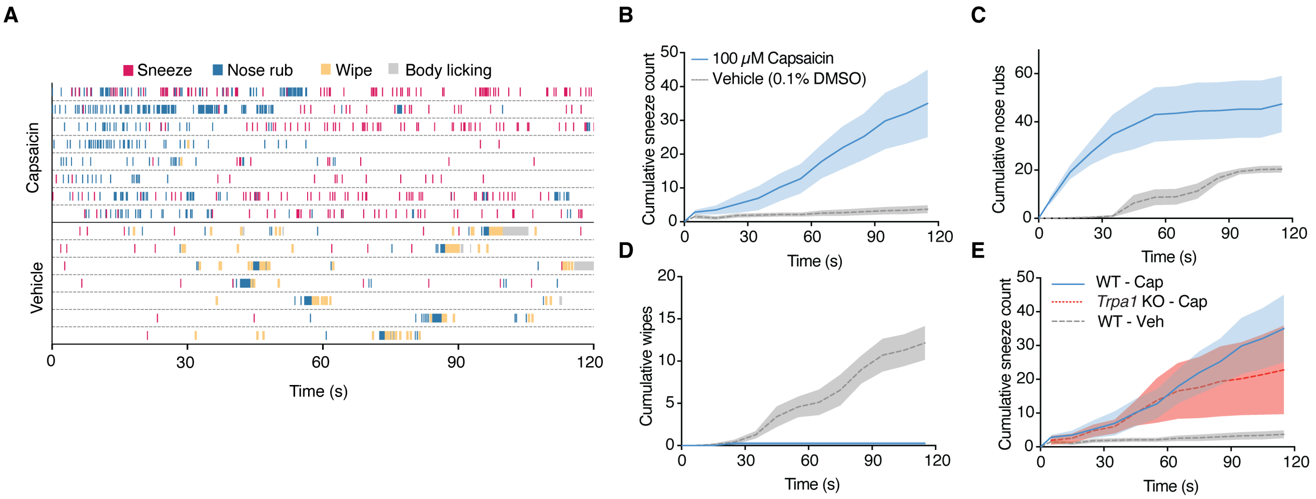


**Figure S5. Capsaicin triggers a variety of intranasal behaviors, related to Figures 1 and 4.**
(A) Raster plot of sneeze, nose rub, wipe, and body-licking events for mice intranasally treated with 100 μM capsaicin or vehicle (0.1% DMSO).
(B–D) Cumulative sneeze count, nose rub count and wipe counts from mice treated with 100 μM capsaicin ($n$ = 8) or vehicle ($n$ = 7).
(E) Cumulative sneeze count of wild-type (WT) and *Trpa1* KO mice treated with 100 μM capsaicin (WT: $n$ = 8, *Trpa1* KO: $n$ = 5) or vehicle (WT: $n$ = 7).
Shading represents mean ± SEM.

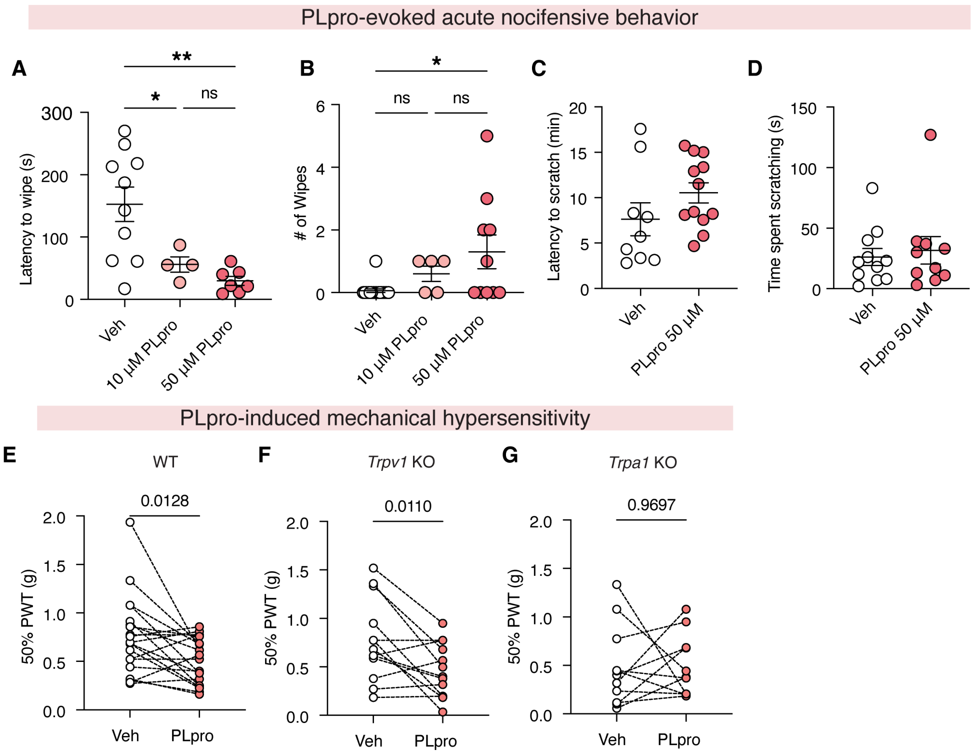


**Figure S6. Intradermal PLpro triggers pain-associated behaviors, related to Figures 1 and 5.**
(A) PLpro elicited a significantly shorter latency to the first wipe. One-way ANOVA: F(2,18) = 9.983, *p* = 0.0012; Tukey's multiple comparisons test: *p* (Veh vs. 10 μM) = 0.0495, *p* (Veh vs. 50 μM) = 0.0028, *p* (10 μM vs. 50 μM) = 0.7890.
(B) PLpro induced a dose-dependent increase in the number of wipes in the first minute. Kruskal-Wallis test: $\chi^2$ = 8.920, *p* = 0.0116; Dunn's multiple comparisons test: *p* (Veh vs. 10 μM) = 0.1340, *p* (Veh vs. 50 μM) = 0.0209, *p* (10 μM vs. 50 μM) > 0.9999. Vehicle (*n* = 17 mice), 10 μM PLpro (*n* = 5 mice), 50 μM PLpro (*n* = 10 mice).
(C) PLpro did not significantly affect the latency or (D) frequency of scratching behavior compared to vehicle over 30 minutes. Vehicle (*n* = 11 mice), 50 μM PLpro (*n* = 10 mice).
(E) WT and (F) *Trpv1* KO mice, but not (G) *Trpa1* KO mice develop mechanical hypersensitivity in the PLpro-injected paw compared to the contralateral vehicle-injected paw 3 hours post-injection. (E) WT: Paired *t*-test, *t* = 2.746, *df* = 19, *p* = 0.0128, *n* = 11 mice. (F) *Trpv1* KO: Paired *t*-test, *t* = 3.001, *df* = 12, *p* = 0.0110, *n* = 13 mice. (G) *Trpa1* KO: paired *t*-test, *t* = 0.039, *df* = 10, *p* = 0.9697, *n* = 11 mice.
Error bars represent mean ± SEM.

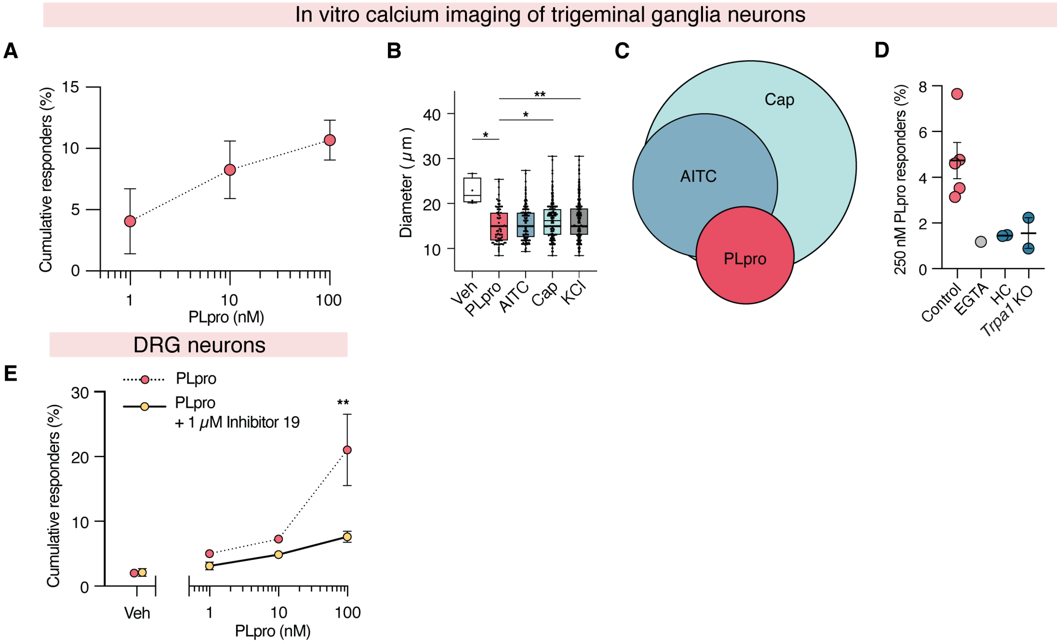


**Figure S7. PLpro activates nociceptors, related to Figure 2.**
(A) PLpro induces dose-dependent activation of neonatal TG neurons in vitro: 1 nM (4.1%), 10 nM (8.3%), and 100 nM (10.7%), *n* = 3 mice.
(B) Diameters of stimulus-responsive TG neurons. PLpro-responsive (combined responders across 1, 10, and 100 nM), neurons were smaller than Veh-, Cap-, and KCl-responsive neurons; Welch's ANOVA: F(4, 22.99) = 6.77, *p* = 0.0009; Dunnett's T3 test versus PLpro: Veh, *p* = 0.0257; AITC, *p* = 0.5442; Cap, *p* = 0.0320; KCl, *p* = 0.0273.
(C) Venn diagram of PLpro-, AITC-, and CAP-responsive TG neurons (*n* = 744 neurons).
(D) Percentage of mouse TG neurons that respond to 250 nM PLpro in control neurons from wild-type C57BL/6N mice in physiological extracellular calcium (2 mM $Ca^{2+}$, 4.7%, *n* = 5) compared to the absence of extracellular calcium (EGTA, 1.2%, *n* = 1), in the presence of a TRPA1 antagonist (HC, 1.4%, *n* = 2), and in neurons from *Trpa1* KO (1.6%, *n* = 2).
(E) Co-application of 1 μM PLpro activity inhibitor 19 significantly reduces the dose-dependent activation of DRG neurons observed with PLpro alone. Two-way repeated measures ANOVA: F(3,6) = 4.891, *p* (Interaction of stimulus and inhibitor) = 0.0473; Šídák's multiple comparisons: *p* (Veh ± inhibitor) > 0.0999, *p* (1 nM ± inhibitor) = 0.9500, *p* (10 nM ± inhibitor) = 0.8920, *p* (100 nM ± inhibitor) = 0.0060, *n* = 2 mice per group.
Error bars represent the mean ± SEM, *n* = biological replicates (animals), with 3-4 wells per animal containing ~100–300 neurons per well.

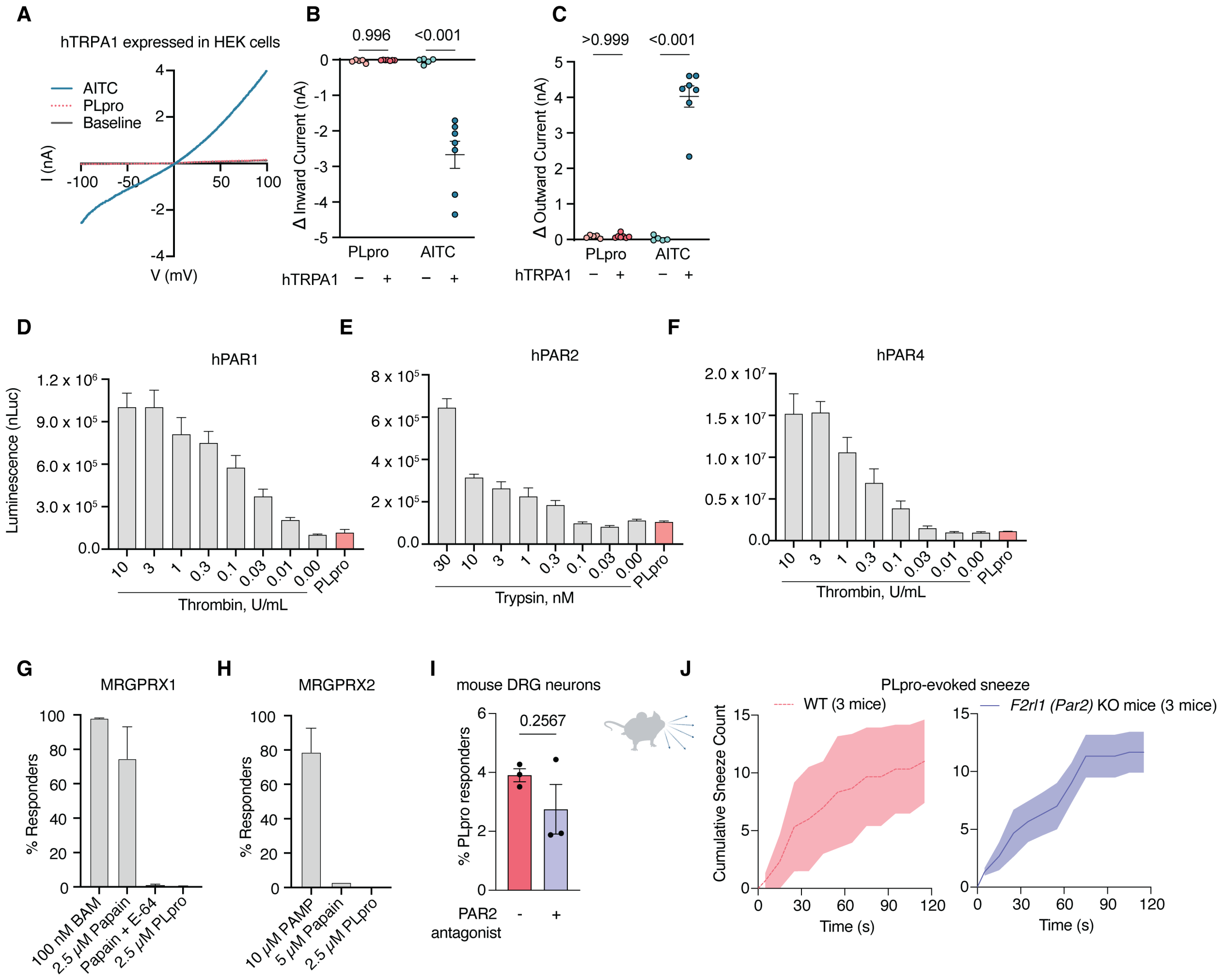


**Figure S8. PLpro does not activate heterologously expressed TRPA1 or other candidate receptors, related to Figure 4.**

(A) Representative current-voltage (I-V) relationships from whole-cell voltage-clamp recordings of human TRPA1-transfected HEK293T cells in the presence of 100 nM PLpro and 100 μM AITC.

(B) Peak inward (−100 mV) currents (baseline-subtracted). One-way ANOVA ($F(3, 20) = 37.7$, $p < 0.001$) with Šídák's multiple comparisons test; *p* (untransfected vs. TRPA1-transfected, PLpro) = 0.996, *p* (untransfected vs. TRPA1-transfected, AITC) < 0.0001.

(C) Peak outward (+100 mV) currents (baseline-subtracted). One-way ANOVA ($F(3, 20) = 137$, $p < 0.001$) with Šídák's multiple comparisons test; *p* (untransfected vs. TRPA1-transfected, PLpro) = >0.999, *p* (untransfected vs. TRPA1-transfected, AITC) < 0.001. PLpro (100 nM) and AITC (100 μM) were each applied for 5 minutes. Each dot represents one cell; data are mean ± SEM (n = number of cells) from three independent experiment days.

(D–F) PAR cleavage assays using nLuc-PAR-eYFP-CHO cells. Cleavage data for human (D) PAR1, (E) PAR2, and (F) PAR4 following treatment with 100 nM PLpro, thrombin (for PAR1 and PAR4), or trypsin (for PAR2).

(G, H) Percentage of Chem-1 cells stably expressing MRGPRX1 or MRGPRX2 that respond to 2.5 μM PLpro, assessed by calcium responses.

(I) Percentage of mouse DRG neurons that respond to 100 nM PLpro in the presence of the PAR2

antagonist, AZ3451, compared to controls (unpaired *t*-test: $t = 1.322$, $df = 4$, $p = 0.2567$, $n = 3$ mice).
(J) *F2rl1* (*Par2*) KO mice exhibit comparable PLpro-evoked sneezing to WT mice, $n = 3$ mice per group. Data are mean ± SEM.

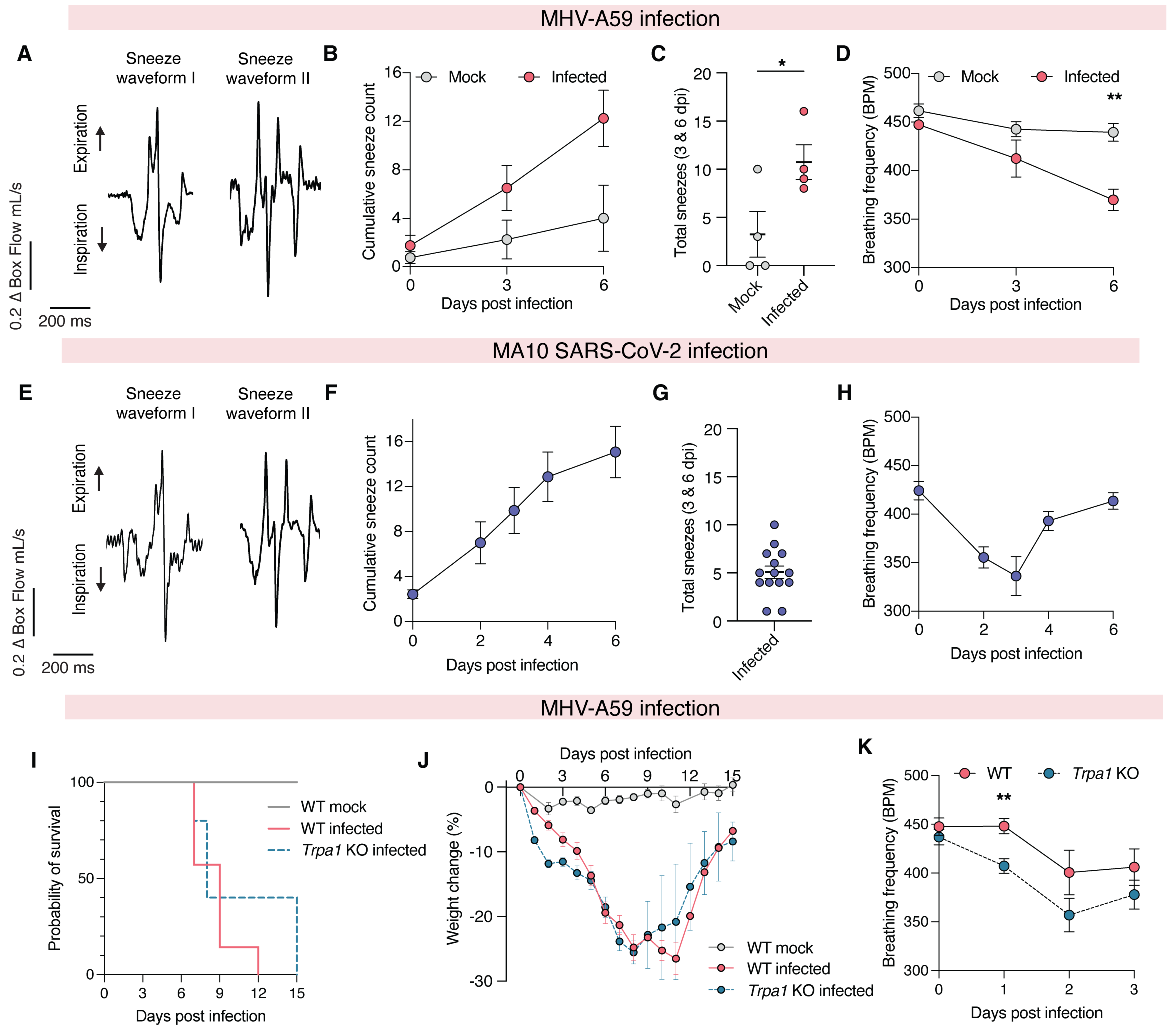


**Figure S9. Sneezing and disease progression during MHV-A59 and mouse-adapted SARS-CoV-2 infection, related to Figure 4.**

(A) Representative box-flow traces of two sneeze waveforms observed following MHV-A59 infection.

(B) Cumulative sneeze counts in mock- and MHV-A59-infected mice at 0, 3, and 6 dpi.

(C) Total sneezes recorded at 3 and 6 dpi in mock- and MHV-A59-infected mice, unpaired *t*-test, $t = 2.529$, $df = 6$, $p = 0.0447$, $n = 4$ mice per group.

(D) Breathing frequency in mock- and MHV-A59-infected mice at 0, 3, and 6 days post-infection (dpi). Two-way repeated-measures ANOVA showed an effect of time ($p = 0.0012$), infection ($p = 0.0138$) or the time × infection interaction ($p = 0.0302$). Šídák's test: Mock- vs. MHV-A59-infected at 6 dpi, $p = 0.0094$; all other time points, ns.

(E) Representative box-flow traces of two sneeze waveforms observed following mouse-adapted (MA10) SARS-CoV-2 infection.

(F) Cumulative sneeze counts in MA10 SARS-CoV-2-infected mice from 0, 2, 3, 4 and 6 dpi.

(G) Total sneezes recorded at 3 and 6 dpi following MA10 SARS-CoV-2 infection.

(H) Breathing frequency in MA10 SARS-CoV-2-infected mice from 0, 2, 3, 4 and 6 dpi, $n = 14$ mice.

(I) Kaplan–Meier survival curves of mock-treated WT and MHV-A59-infected WT and Trpa1 KO mice (25,000 PFU). Survival did not differ between infected genotypes, log-rank Mantel–Cox test: $\chi^2 = 1.011$, $p = 0.3146$.

(J) Percentage change in body weight following mock infection of WT (*n* = 5 mice) or MHV-A59 infection of WT (*n* = 8 mice) and *Trpa1* KO mice (*n* = 5 mice).

(K) Breathing frequency in MHV-A59-infected WT and *Trpa1* KO mice from 0 to 3 dpi. Two-way repeated-measures ANOVA showed an effect of time ($p = 0.0002$), but not genotype ($p = 0.0702$) or the time and genotype interaction ($p = 0.3562$). Šídák's test: WT versus *Trpa1* KO at 1 dpi, $p = 0.0082$; all other time points, ns.

Data are mean ± SEM. *$p < 0.05$ and **$p < 0.01$.

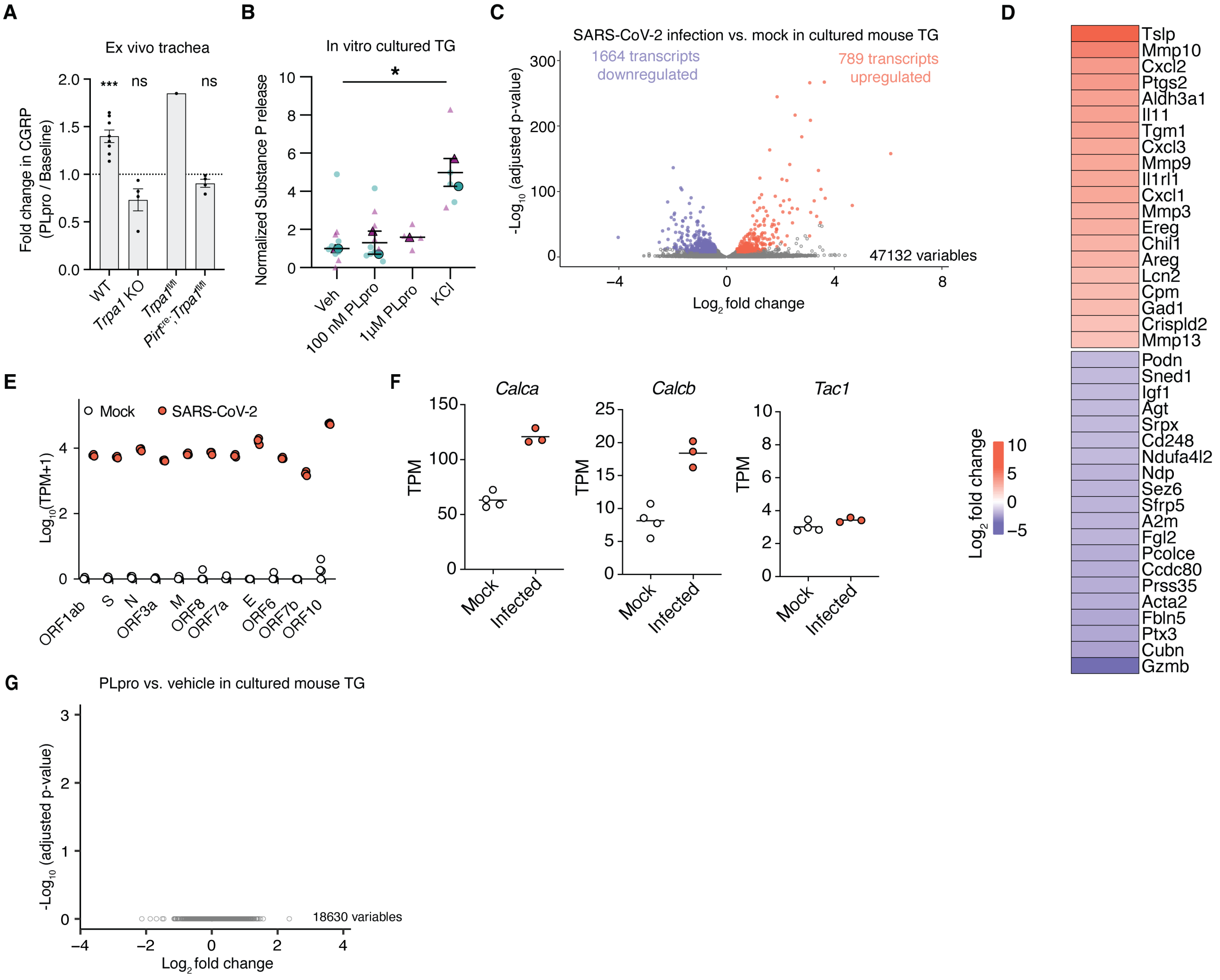


**Figure S10. PLpro evokes CGRP but not substance P release, whereas SARS-CoV-2 infection alters gene expression in cultured mouse trigeminal ganglia, related to Figure 5.**

(A) PLpro-evoked CGRP release from ex vivo tracheal preparations from WT, *Trpa1* KO, *Trpa1*$^{fl/fl}$ and *Pirt*$^{cre}$;*Trpa1*$^{fl/fl}$ mice expressed as fold over baseline. The dotted line denotes baseline release (fold change = 1). PLpro increased CGRP release above baseline in WT trachea (one-sample *t*-test: $t$ = 6.040, $df$ = 7, $p$ = 0.0005; $n$ = 8), but not in *Trpa1* KO ($t$ = 2.322, $df$ = 3, $p$ = 0.1029; $n$ = 4) or *Pirt*$^{Cre}$;*Trpa1*$^{fl/fl}$ trachea ($t$ = 2.222, $df$ = 3, $p$ = 0.1129; $n$ = 4). No statistical test was performed for the *Trpa1*$^{fl/fl}$ control ($n$ = 1). Data are mean ± SEM.

(B) PLpro does not elicit the release of Substance P from cultured trigeminal ganglia (TG). One-way ANOVA: $F(3,3)$ = 11.44, $p$ = 0.0377; Dunnett's post hoc test: $p$ (Veh vs. 100 nM PLpro) = 0.9577, $p$ (Veh vs. 1 μM PLpro) = 0.8682, $p$ (Veh vs. KCl) = 0.0303, Vehicle ($n$ = 2 biological replicates), 100 nM PLpro ($n$ = 2 biological replicate), 1 μM PLpro ($n$ = 1 biological replicate), KCl ($n$ = 2 biological replicates). Each biological replicate contains pooled TG, with 2-6 wells per biological replicate.

(C) SARS-CoV-2 infection for 48 hours upregulates 789 transcripts and downregulates 1,664 transcripts compared to mock infection. Differentially expressed genes were identified using the following criteria: base mean > 152, absolute $\log_2$ fold change > 0.37, and adjusted p-value < 0.01. SARS-CoV-2-infected ($n$ = 3 wells) and mock-infected ($n$ = 4 wells), $n$ represents wells from pooled TG from 4 mice.

(D) Heatmap of the 20 most highly upregulated and downregulated transcripts ($\log_2$ fold change of

SARS-CoV-2 infection vs. mock).
(E) SARS-CoV-2 viral transcript abundance (transcript per million [TPM]) in cultured TG 48 hours post-infection.
(F) *Calca* and *Calcb* but not *Tac1* transcripts are significantly upregulated by SARS-CoV-2 infection. Adjusted $p$ (*Calca)* = $9.05 \times 10^{-21}$, adjusted $p$ (*Calcb)* = $5.52 \times 10^{-9}$; adjusted $p$ (*Tac1)* = 0.991.
(G) Treatment with 2 μM PLpro for 14-16 h did not alter transcript expression in cultured TG compared with vehicle (PLpro, $n$ = 6 biological replicates; vehicle, $n$ = 5 biological replicates). Each replicate contained TG pooled from 4 mice.
Data are mean ± SEM.

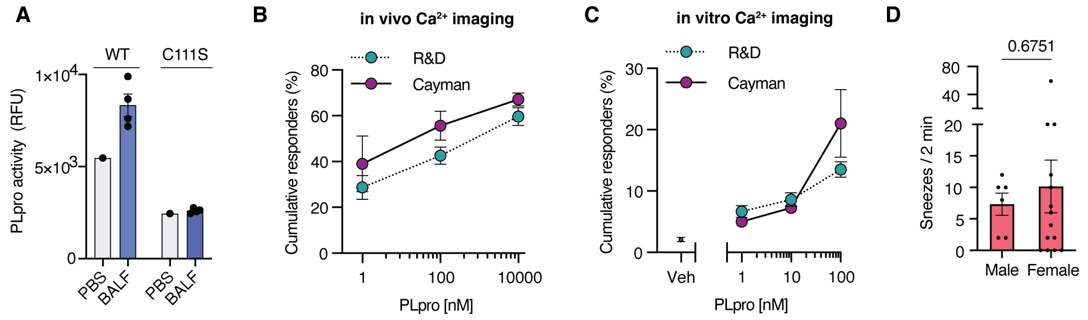


**Figure S11. Recombinant PLpro displays proteolytic activity in bronchoalveolar lavage fluid and induces comparable neuronal responses across suppliers, with no sex difference in PLpro-induced sneezing, related to Figure 1 and 2.**

(A) Recombinant wild-type (WT) or catalytically inactive (C111S) PLpro was preincubated with PBS or bronchoalveolar lavage fluid (BALF) and proteolytic activity was measured after 1 hour, by cleavage of the fluorogenic peptide substrate Z-RLRGG-AMC, reported as relative fluorescence units (RFU). BALF was collected by intratracheal PBS lavage from 4 individual mice; dots represent the mean of two technical replicates per animal. PBS wells contained recombinant PLpro alone in buffer.
(B) Cumulative percentage of trigeminal neurons responding to PLpro from R&D and Cayman in vivo. Two-way ANOVA**:** $F(2,8) = 10.18$, $p$ (R&D vs. Cayman) = 0.2451; Šídák's multiple comparisons**:** $p$ (1 nM) = 0.6320, $p$ (100 nM) = 0.4480, $p$ (10 µM) = 0.8183, $n$ = 3 mice per group.
(C) PLpro from both suppliers induces dose-dependent activation of DRG neurons in vitro. Two-way ANOVA**:** $F(1,7) = 7.140$, $p$ (R&D vs. Cayman) = 0.5455; Šídák's multiple comparisons**:** $p$ (veh) = 0.9951, $p$ (1 nM) = 0.5219, $p$ (10 nM) = 0.6979, $p$ (100 nM) = 0.8668, R&D ($n$ = 7 mice), Cayman ($n$ = 2 mice). Error bars represent the mean ± SEM, $n$ = biological replicates (animals), with 3–4 wells per animal containing ~100–300 neurons per well.
(D) Number of sneezes over 2 minutes in male compared with female mice (unpaired t-test: $t = 0.4261$, df = 18, $p = 0.6751$, n = 6 male mice and 14 female mice).

| Protease | Supplier | Preparation | Tag | Endotoxin (EU/mL per nanomolar) | Purity (Vendor-reported) |
|---|---|---|---|---|---|
| SARS-CoV-2 $PLpro_{WT}$ | Cayman #31817 | Lyophilized from sterile 20 mM Tris, pH 7.4, with 500 mM sodium chloride | N-terminal His-tag | $1.00 \times 10^{-3} \pm 4.42 \times 10^{-4}$ | ≥90% estimated by SDS-PAGE |
| SARS-CoV-2 $PLpro_{WT}$ | R&D #E-611 | In 50 mM HEPES, 300 mM NaCl, 1 mM TCEP, 10% (v/v) Glycerol, pH 8.0 | N-terminal GST-tag | $1.80 \times 10^{-1} \pm 4.51 \times 10^{-4}$ | >90%, by SDS-PAGE under reducing conditions and visualized by Colloidal Coomassie® Blue stain. |
| SARS-CoV-2 $PLpro_{C111S}$ | R&D custom order | In 50 mM HEPES, 300 mM NaCl, 1 mM TCEP, 10% (v/v) Glycerol, pH 8.0 | N-terminal GST-tag | $6.08 \times 10^{-2} \pm 5.16 \times 10^{-4}$ | 87.4% by SDS-PAGE under reducing conditions and visualized by Colloidal Coomassie® Blue stain. |
| SARS-CoV PLpro | R&D #E-610 | In 50 mM HEPES, 100 mM NaCl, 2 mM TCEP, pH 7.5 | N-terminal 6-His tag | $2.58 \times 10^{-4} \pm 1.38 \times 10^{-4}$ | >95%, by SDS-PAGE under reducing conditions and visualized by Colloidal Coomassie® Blue stain. |
| MERS-CoV PLpro | R&D #E-609 | In 50 mM HEPES, 200 mM NaCl, 1 mM DTT, 10% (v/v) Glycerol, pH 7.5 | N-terminal 6-His tag | $1.49 \times 10^{-4} \pm 1.72 \times 10^{-4}$ | >90%, by SDS-PAGE under reducing conditions and visualized by Colloidal Coomassie® Blue stain. |
| SARS-CoV-2 Mpro | R&D #E-720 | In 50 mM HEPES, 150 mM NaCl, 1 mM DTT, 1 mM EDTA, pH 7.5 | None | $5.42 \times 10^{-1} \pm 5.18 \times 10^{-2}$ | >95%, by SDS-PAGE under reducing conditions and visualized by Colloidal Coomassie® Blue stain. |
| Vehicle | -- | 50 mM HEPES, 300 mM NaCl, 1 mM TCEP, 10% (v/v) Glycerol, pH 8.0 | | $4.00 \times 10^{-4}$ | |

**Table S1**. **Recombinant protease properties, related to Figure 3.**
Endotoxin measurements were determined by the LAL assay and displayed as mean ± SEM. All other data are vendor-provided.
